# A validated panel of commercial antibodies for reliable detection of FET proteins

**DOI:** 10.1101/2025.11.19.689317

**Authors:** Sara Tacconelli, Laura Odemwingie, Stefan Reber, Sergio Di Pancrazio, Tatjana I Zoller, Luca Biasetti, Claire Troakes, Marc-David Ruepp, Caroline Vance

## Abstract

The FET protein family comprises the highly conserved RNA-binding proteins FUS, EWS and TAF15 which are implicated in RNA metabolism and neurodegenerative diseases such as ALS and FTLD. Despite their structural similarity, reliable detection of individual FET proteins remains challenging due to antibody cross-reactivity and inconsistent localisation patterns reported in the literature. To address this, we systematically evaluated 23 commercially available antibodies for specificity and performance across western blotting, immunocytochemistry, and immunohistochemistry. Using single and double shRNA knockdowns in HeLa cells, we confirmed target engagement and identified antibodies which were specific as well as those with significant cross-reactivity. Immunofluorescence in rat primary neurons and HeLa cells showed antibody and cell type dependent variations in nuclear and cytoplasmic localisation; from these, we identified a subset that demonstrated high-quality, region-specific staining in postmortem human brain tissue. Our findings highlight substantial variability in antibody performance and underscore the need for rigorous validation to ensure reproducibility in FET protein research. We present a validated panel of antibodies suitable for diverse applications, providing a critical resource for studies of FET protein biology and pathology.

## Introduction

The RNA-binding proteins Fused in Sarcoma (FUS), Ewing Sarcoma (EWS) and TATA-box binding protein-associated factor 15 (TAF15) are ubiquitously expressed, highly conserved, nucleo-cytoplasmic shuttling RNA binding proteins, that make up the so-called FET family of proteins (Schwartz et al., 2015). Their domain organisation is highly similar, consisting of an N-terminal domain of low complexity, followed by an RNA recognition motif (RRM) and Zinc Finger domain (ZnF), each flanked by arginine–glycine–glycine (RGG)-rich domains, and a C-terminal PY-Nuclear localisation signal (NLS) (Figure 1). The PY-NLS is recognised by Transportin-1 (TNPO1), which mediates their import into the nucleus (Lee et al., 2006; Marko et al., 2012). In 2009, mutations in the FUS gene were identified that cause mislocalisation and aggregation of FUS in the cytoplasm of neurons and glial cells resulting in the development of Amyotrophic Lateral Sclerosis (Kwiatkowski et al., 2009; Vance et al., 2009). In the same year, FUS was also found to be present in pathological cytoplasmic inclusions in a subset of Frontotemporal Lobar Degeneration (FTLD) cases (Neumann et al., 2009). However, it has since been discovered that in these FTLD cases FUS co-aggregates not only with TNPO1, but also with the other members of the FET family with TAF15 forming amyloid filaments (Brelstaff et al., 2011; Davidson et al., 2013; Gami-Patel et al., 2016; Neumann et al., 2011; Neumann et al., 2012; Tetter et al., 2024; Van Langenhove et al., 2010).

**Figure 1:**
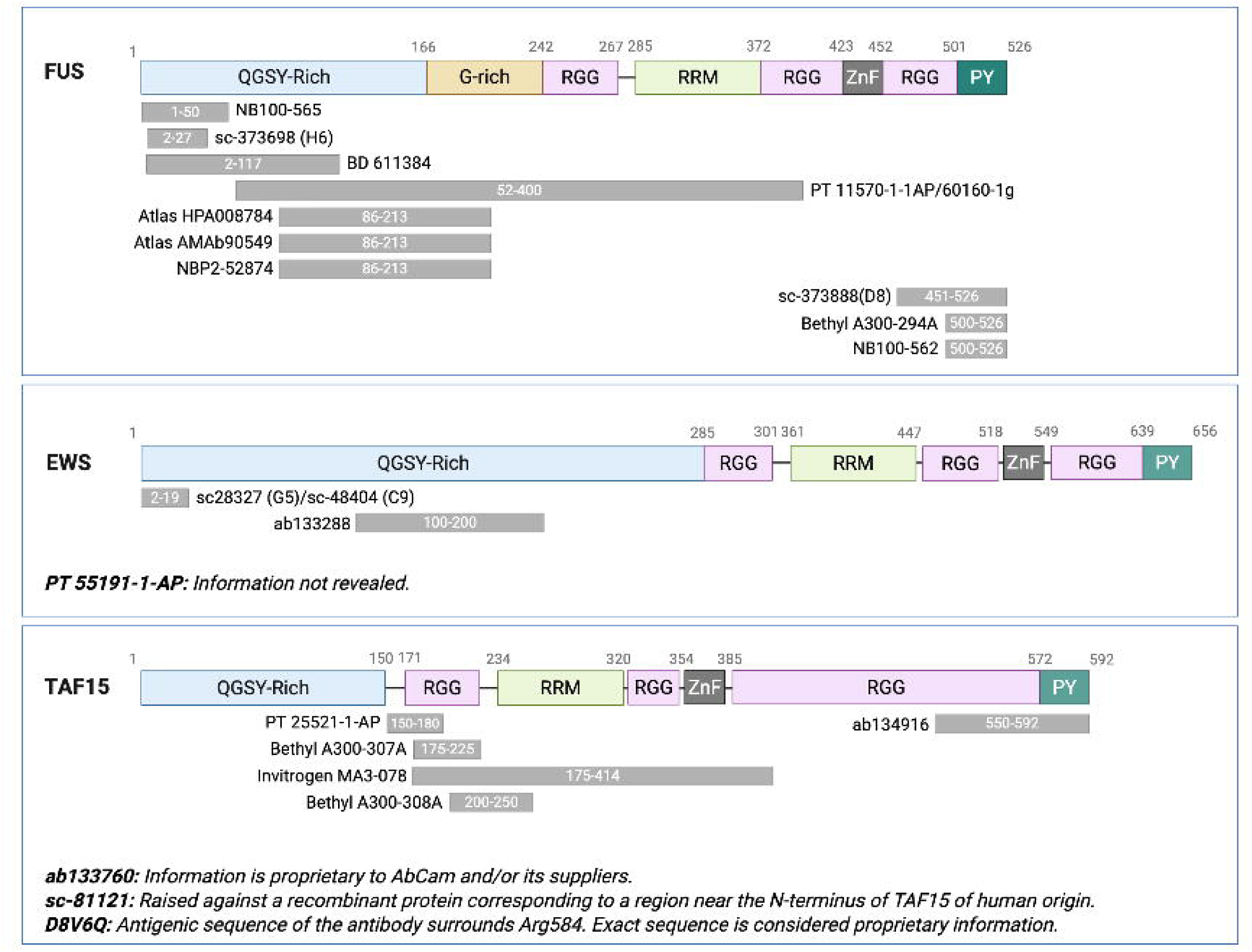
Schematic of FET family protein architecture. Epitopes of the different anti-FUS, anti-EWS and anti-TAF15 antibodies have been indicated where they are known. Domains are labelled for their function/make up: low complexity domain (QGSY Rich) RNA recognition motif (RRM) and Zinc Finger domain (ZnF), each flanked by arginine-glycine–glycine (RGG)-rich domains, and a C-terminal PY-Nuclear localisation signal (NLS).

In the absence of disease, the FET proteins have mainly been described as predominantly nuclear in line with their involvement in diverse steps of RNA metabolism ranging from transcription to pre-mRNA splicing (Schwartz et al., 2015; Tan and Manley, 2009) with FUS and TAF15 displaying additional cytoplasmic localisation (Andersson et al., 2008; Blechingberg et al., 2012). However, more and more reports suggest additional functions of FET-family members outside the nucleus, such as at the synapse or in the context of local translation (Salam et al., 2021; So et al., 2018; Yasuda et al., 2013). Intriguingly, depending on the antibodies and methodology used to detect the family members, different nucleo/cytoplasmic localisation ratios have been reported as well as epitope-specific outcomes (Tsai et al., 2022).

There is a high degree of conservation among FET protein family members in humans, with sequence identities of 46.9% (TAF15-EWS), 54.3% (FUS-EWS), and 60.3% (FUS-TAF15). Whilst invertebrates and plants have only a single FET protein, all vertebrates have three which are highly conserved across species suggesting distinct specialised roles for each family member (Shwartz et al., 2015). Due to the high conservation and reported differences in localisation outcomes in published studies, we questioned whether this might not only stem from epitope location and availability but also from antibody specificity and cross-reactivity. Given the critical reliance on antibodies for studying both the physiological functions of individual FET proteins and their roles in disease - especially in the context of the ongoing antibody characterisation crisis (Kahn et al., 2024) - we systematically evaluated 23 commercially available FET antibodies for target engagement and cross-reactivity. Here, we present a validated panel of antibodies specific for each FET member, suitable for use in western blotting, immunofluorescence, and immunohistochemistry (IHC).

## Results

### The FET proteins show a high degree of similarity

The FET proteins show a high degree of conservation with similar domain architectures (Figure 1) each containing an N-terminal low complexity domain, 3 RGG domains, an RNA recognition motif (RRM), a Zinc finger domain (ZnF) as well as a C-terminal nuclear localisation signal (NLS). Twenty-three different antibodies against the FET proteins were tested, and where epitope information was available, the epitopes were mapped onto the protein sequence (Figure 1). While for both FUS and TAF15, we included antibodies raised against various regions and domains, for EWS, only antibodies raised against the N-terminus were tested due to availability.

### FET antibodies cross react with other members of the family

To determine whether each antibody can discriminate its target FET protein from its closely related family members a single and double knockdown assay was performed in HeLa cells. Cells were transfected with shRNAs against FUS, EWS or TAF15 and the combinations of FUS+EWS, EWS+TAF15 and FUS+TAF15. Knockdown was confirmed with qPCR showing efficacy for each shRNA panel (Figure S1). FUS mRNA levels in the FUS only KD, FUS+EWS KD and FUS+TAF15 KD were significantly lower compared to control (p<0.01). EWS mRNA levels only show a significant decrease in EWS only KD, FUS+EWS KD and EWS + TAF15 KD compared to control (p<0.01). Similarly, TAF15 mRNA levels only significantly decrease in TAF15 only KD, FUS + TAF15 KD and EWS+TAF15 KD compared to control (p<0.05). We also observed evidence of potential cross-knockdown, as qPCR for FUS mRNA revealed a reduction in FUS expression in TAF15 shRNA treated samples. This suggests that one or both of the TAF15-targeting shRNAs may also partially target FUS mRNA. Following on, antibodies raised against the FET proteins were validated by western blot using lysates from the shRNA transfected cells. Antibodies were selected from a range of commercial companies attempting to identify those raised in both mouse and rabbit that were specific for each FET family member (Table 1). Whilst this is not a complete set of commercially available antibodies, it encompasses those used routinely in publications from within the field and our own labs.

| Antigen | Company | Code | Host species | Optimised dilutions |  |  | Specific |
| --- | --- | --- | --- | --- | --- | --- | --- |
|  |  |  |  | WB | ICC | IHC |  |
| FUS | BD | 611384 | Mouse | 1:1000 | 1:500 |  | Yes |
| FUS | Protein tech | 60160-1-Ig | Mouse | 1:1000 | 1:200 |  | No |
| FUS | Santa Cruz | sc-373698 (H6) | Mouse | 1:500 | 1:200 |  | No |
| FUS | Santa Cruz | sc-373888 (D8) | Mouse | 1:100 | 1:200 |  | Yes |
| FUS | Sigma (Atlas) | AMAb90549 | Mouse | 1:1000 | 1:200 |  | Yes |
| FUS | Novus | NBP2-52874 | Mouse | 1:1000 | 1:200 |  | Yes |
| FUS | Protein tech | 11570-1-AP | Rabbit | 1:200 | 1:200 |  | Yes |
| FUS | Bethyl | A300-294A | Rabbit | 1:1000 | 1:500 |  | No |
| FUS | Novus | NB100-565 | Rabbit | 1:1000 | 1:500 |  | Yes |
| FUS | Novus | NB100-562 | Rabbit | 1:1000 | 1:200 |  | No |
| FUS | Sigma (Atlas) | HPA008784 | Rabbit | 1:1000 | 1:1000 | 1:100 | Yes |
| EWS | Santa Cruz | sc-48404 (C9) | Mouse | 1:500 | 1:200 |  | Yes |
| EWS | Santa Cruz | sc-28327 (G5) | Mouse | 1:500 | 1:200 |  | Yes |
| EWS | Abcam | ab133288 | Rabbit | 1:500 | 1:200 | 1:100 | Yes |
| EWS | Protein tech | 55191-1-AP | Rabbit | 1:1000 | 1:200 |  | No |
| TAF15 | Invitrogen | MA3-078 (8TA-2B10) | Mouse | 1:1000 | 1:200 |  | Yes |
| TAF15 | Santa Cruz | sc-81121 | Mouse | 1:200 | 1:200 |  | Yes |
| TAF15 | Abcam | ab133760 | Rabbit | 1:500 | 1:250 |  | Yes |
| TAF15 | Abcam | ab134916 | Rabbit | 1:500 | 1:250 | 1:100 | Yes |
| TAF15 | Bethyl | A300 307A | Rabbit | 1:500 | 1:200 |  | Yes |
| TAF15 | Bethyl | A300 308A | Rabbit | 1:500 | 1:200 |  | Yes |
| TAF15 | Protein tech | 25521-1-AP | Rabbit | 1:200 | 1:100 |  | No |
| TAF15 | Cell Signalling | 28409 (D8V6Q) | Rabbit | 1:1000 | 1:500 |  | Yes |

Eleven different antibodies from six different providers against FUS were tested (Table 1) and of these, seven showed clear specificity for FUS with a band identified at ∼70KDa which was reduced in the FUS shRNA treated samples with no other bands observed (Figure 2 with quantification graphs in Figure S2 and full blots in Figure S3). Four different mouse antibodies, AMAb90549 from Sigma, sc-373888 (D8) from Santa Cruz, 611384 by BD Biosciences and NBP2-52874 by Novus, all showed a single band at the expected molecular weight, the expression of which was lowered when shRNA FUS was expressed (Figure 2A-D, Figure S2A-D, Figure S3A-D: control vs FUS KD p<0.0001, control vs FUS+EWS KD p=0.0046, control vs FUS+TAF15 KD p=0.016; control vs FUS KD p<0.0001, control vs FUS+EWS KD p<0.0001, control vs FUS+TAF15 KD p<0.0001; control vs FUS KD p=0.0131, control vs FUS/EWS KD p=0.0076, control vs FUS/TAF15 KD p=0.0106; control vs FUS KD p<0.05, control vs FUS+EWS KD p<0.05; control vs FUS+TAF15KD p=0.0.05). The only additional significant change in FUS expression was that for sc-373888, FUS was also affected by TAF15 knockdown in line with the FUS mRNA qPCR results (control vs TAF15 KD, p=0.003; control vs EWS+TAF15 KD, p=0.0001). Three rabbit anti-FUS antibodies, HPA008784 antibody by Atlas (Sigma-Aldrich), Novus NB100-565 and 11570-1-AP by ProteinTech, all showed one clear band, at around 70kDa, the intensity of which was affected in the samples depleted from FUS (Figure 2E-G, Figure S2E-G, Figure S3E-G: control vs FUS KD p=0.0017, control vs FUS+EWS KD p=0.0007, control vs FUS+TAF15 KD p=0.0032; control vs FUS KD p=0.0845; control vs FUS+EWS KD p=0.0742; control vs FUS+TAF15 KD p=0.1872; control vs FUS KD p=0.0058; control vs FUS+EWS KD p=0.0046; control vs FUS+TAF15KD p=0.016) although this did not reach significance for NB100-565.

**Figure 2:**
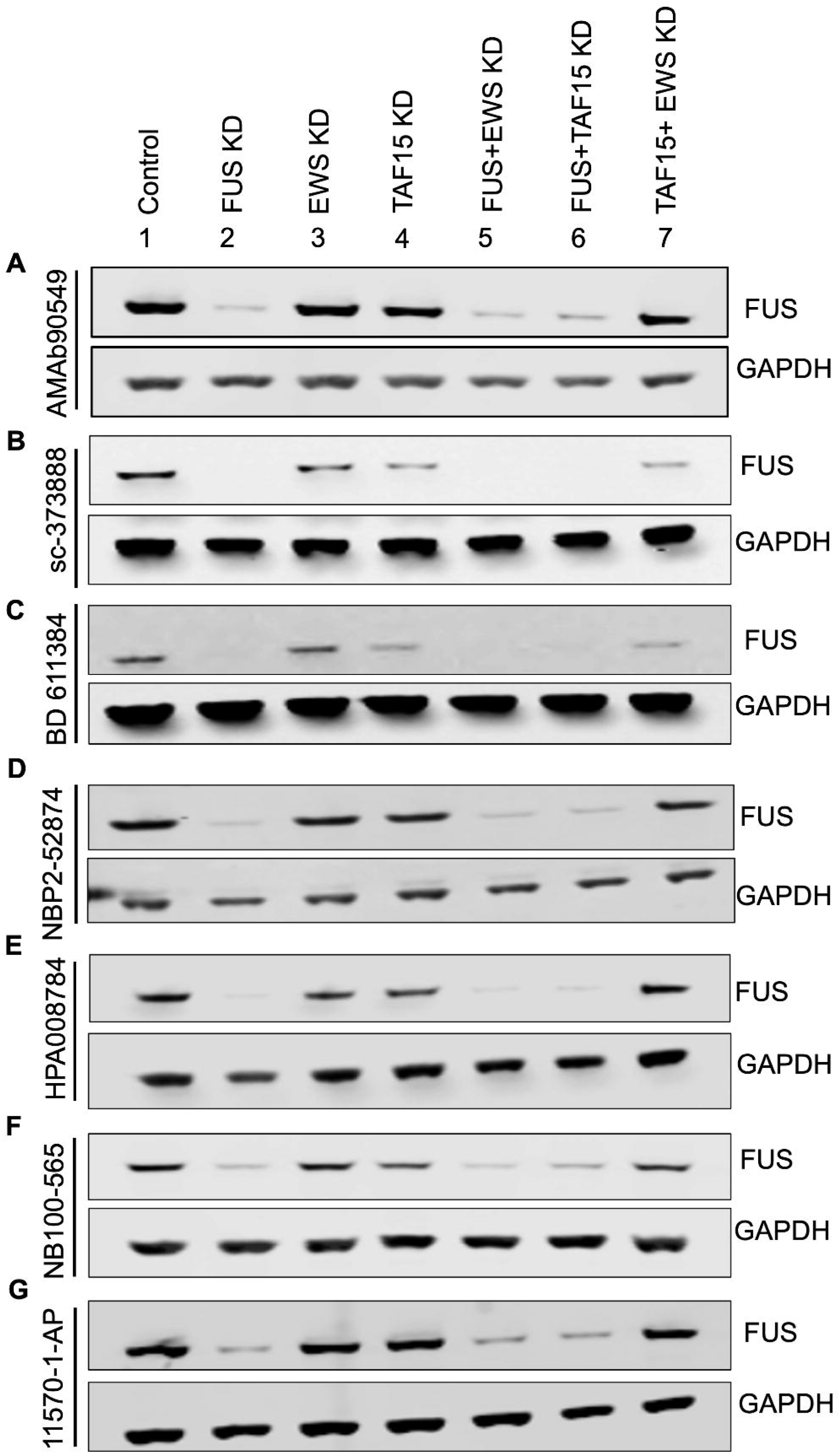
Specific anti-FUS antibodies confirmed by single/double FET shRNA HeLa knockdowns. Western blot validation of seven commercially available anti-FUS antibodies showed a single band at ∼70 kDa corresponding to FUS. All antibodies show a decrease of FUS protein levels in FUS only KD (Lane 2), FUS + EWS KD (Lane 5) and FUS + TAF15 KD (Lane 6). All bands were normalised to GAPDH loading control (37 kDa) and ratios of FET single/double knockdowns were then compared to control and significance determined by performing One-Way ANOVA with post-hoc Tukey’s multiple comparisons test (See Figure S2 for quantifications).

Four of the anti-FUS antibodies showed additional bands at different molecular weights (Figure 3, Figure S4 & Figure S5). Two of them, the rabbit Bethyl A300-294A and the mouse sc373698 antibodies stained for two clear bands, one at ∼80kDa and one at ∼70kDa (Figure 3A&B, Figure S4A&B & Figure S5A&B). The lower band was significantly reduced when FUS was knocked down, and so was identified as FUS (control vs FUS KD p<0.0001, control vs FUS+EWS KD p<0.0001, control vs FUS+TAF15 KD p<0.0001; control vs FUS KD p=0.0058, controls vs FUS+EWS KD p=0.0046, control vs FUS+TAF15 KD, p=0.016) and the higher one was reduced upon EWS depletion suggesting the antibody cross reacts with EWS (control vs EWS KD p<0.0001, control vs FUS+EWS KD p<0.0001, control vs EWS+TAF15 KD p<0.0001; control vs EWS KD p=0.157, control vs FUS+EWS KD p=0.1132, control vs EWS+TAF15 KD p=0.1136). The mouse anti-FUS antibody from Protein Tech (60160-1-Ig) displayed visible bands at seven different molecular weights (Figure 3C, Figure S4C and Figure S5C). Whilst the identity of four of these bands remains elusive (Bands A, B, C & D at 90kDa, 55kDa, 48kDa and 42kDa respectively), there was a band at ∼70kDa which was reduced upon FUS knockdown indicating it does bind to FUS (control vs FUS KD p<0.01, control vs FUS+EWS KD p<0.01, control vs FUS+TAF15 KD p<0.05). However, this antibody seems to cross-react with EWS as shown by the band at ∼80kDa that is reduced upon EWS depletion (control vs EWS KD p=0.014, control vs FUS+EWS KD p=0.0012, control vs EWS+TAF15 KD p=0.0016). There is also a faint band that is reduced when TAF15 is knocked down suggesting it is TAF15 (control vs TAF15 KD p=0.0009, control vs FUS+TAF15 p=0.0018, control vs EWS+TAF15 p=0.009). This suggests that this antibody is has a high degree of cross-reactivity and possibly picks up additional or fragmented FET protein products. The final antibody we tested that was raised against FUS was NB100-562, resulted in four different bands, two at around 80kDa, one at ∼70kDa and one at ∼35kDa of which two appeared to correspond to FET proteins (Figure 3D, Figure S4D, Figure S5D). The intensity of the band at 70kDa was reduced upon FUS depletion confirming this antibody recognises FUS (control vs FUS KD p=0.004, controls vs FUS+EWS KD p=0.0033, control vs FUS+TAF15 KD p=0.0078) while the higher band at ∼80kDa was identified as EWS (control vs EWS KD p=0.0488, control vs FUS+EWS KD p=0.0451, control vs EWS+TAF15 KD p=0.0574).

**Figure 3:**
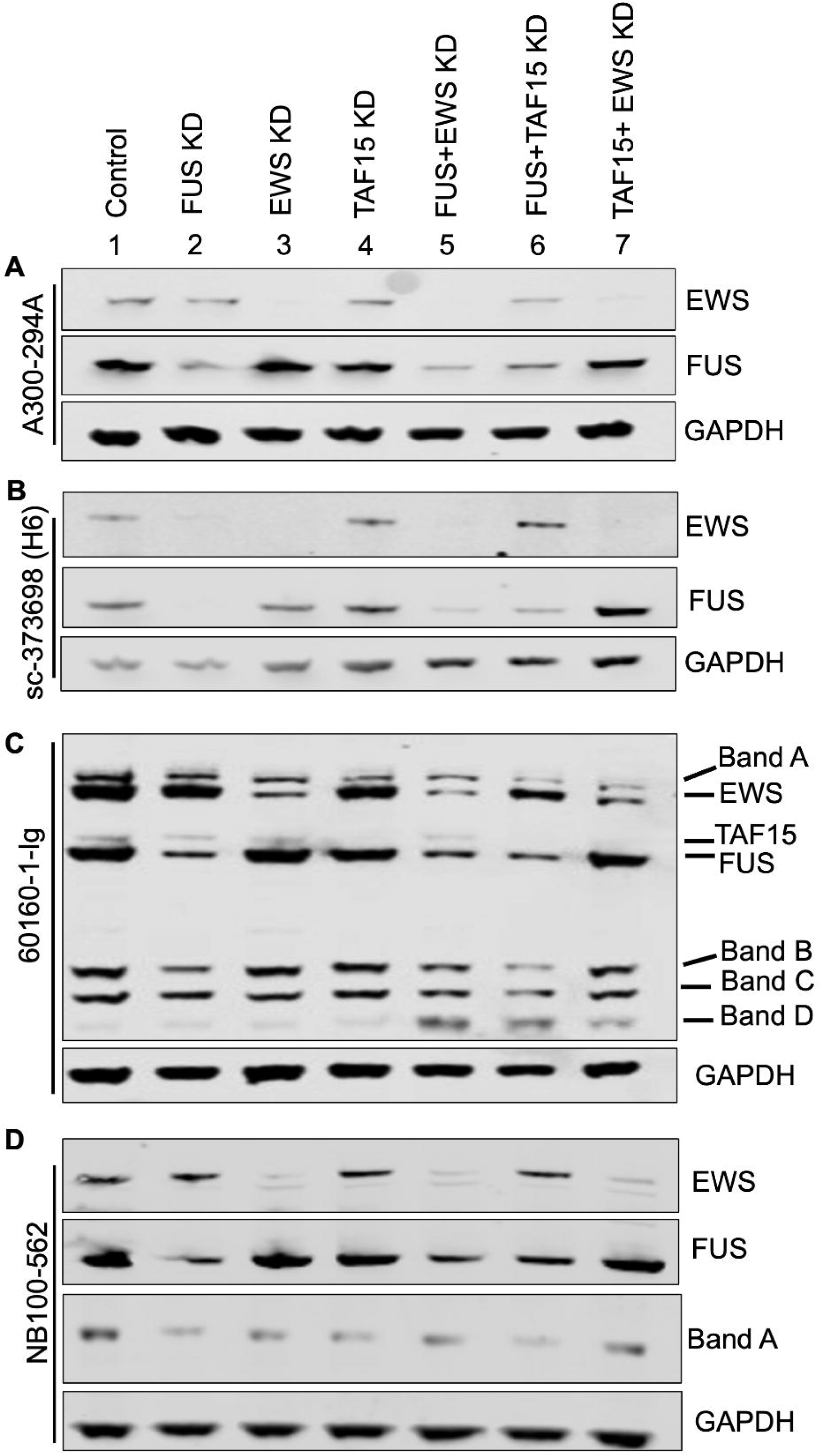
Cross-reactivity of some anti-FUS antibodies confirmed by single/double FET shRNA HeLa knockdowns. Western blot validation of four commercially available anti-FUS antibodies showed a single band at ∼70 kDa corresponding to FUS. All antibodies show a decrease of FUS protein levels in FUS only KD (Lane 2), FUS + EWS KD (Lane 5) and FUS+TAF15 KD (Lane 6). However, all four showed additional bands identified as EWS and/or TAF15 as well as those that don’t appear to recognise any FET protein. All bands were normalised to GAPDH loading control (37 kDa) and ratios of FET single/double knockdowns were then compared to control and significance determined by performing One-Way ANOVA with post-hoc Tukey’s multiple comparisons test (See Figure S4 for quantifications).

We tested four different antibodies raised against EWS (Table 1, Figure 4, Figure S6 for quantifications and Figure S7 for full blots). Three of them, anti-EWS sc-48404 and sc-28237 from Santa Cruz and Abcam ab133288 produced robust bands that were strongly reduced when EWS was knocked-down (Figure 4A-C, Figure S6A-C, Figure S7A-C, control vs EWS KD p<0.0001, control vs FUS+EWS KD p<0.0001, control vs EWS+TAF15 KD p<0.0001; control vs EWS KD p=0.1360, control vs FUS+EWS KD p=0.1170, control vs EWS+TAF15 KD p=0.1434; control vs EWS KD p<0.0001, control vs FUS+EWS KD p<0.000, control vs EWS+TAF15 KD, p<0.0001) though this only reached significance for sc-48404 and ab133288. Further, EWS on the ab133788 showed a significant reduction in EWS signal when FUS and TAF15 levels were lowered by shRNA (control vs FUS+TAF15 KD, p=0.0139). When evaluating the rabbit ProteinTech anti- EWS 55191-1-AP, multiple faint bands were detected. However, one prominent band was consistently observed that was significantly reduced in the samples where EWS levels was knocked down (Figure 4D, Figure S6D, Figures S7D; control vs EWS KD p=0.0002; control vs FUS+EWS KD p=0.0002; control vs EWS+TAF15 KD p=0.0002), but also slightly reduced when both FUS and TAF15 were knocked down (control vs FUS+TAF15, p=0335).

**Figure 4:**
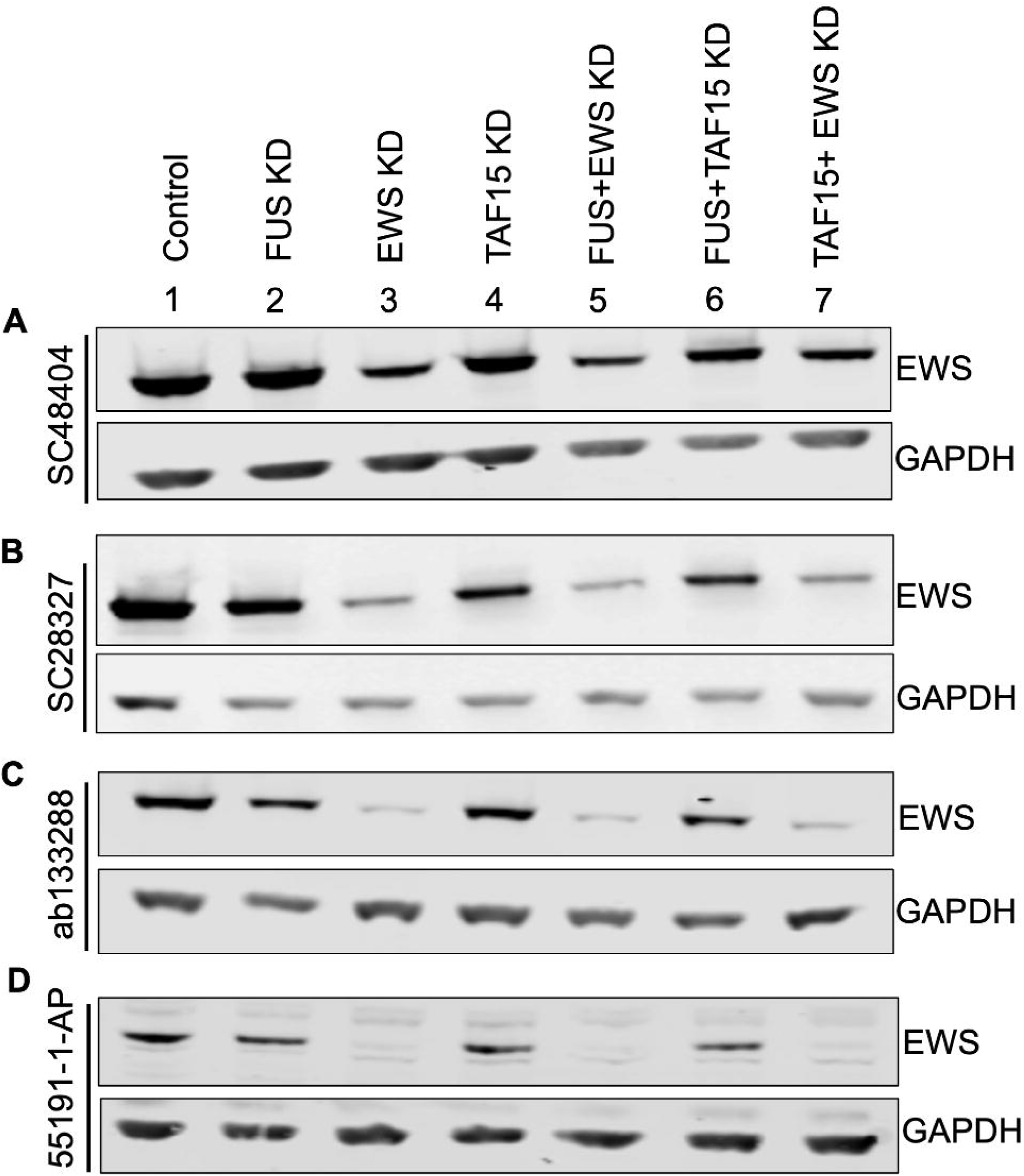
Specific anti-EWS antibodies confirmed by single/double FET shRNA HeLa knockdowns. Western blot validation of seven commercially available anti-EWS antibodies showed a single band at ∼80 kDa corresponding to EWS. All antibodies show a decrease of EWS protein levels in EWS only KD (Lane 3), FUS + EWS KD (Lane 5) and TAF15 +EWS KD (Lane 7). All bands were normalised to GAPDH loading control (37 kDa) and ratios of FET single/double knockdowns were then compared to control and significance determined by performing One-Way ANOVA with post-hoc Tukey’s multiple comparisons test (See Figure S6 for quantifications).

Eight different TAF15 antibodies were tested in our knockdown system (Table 1). Across virtually all antibodies, two immunoreactive bands were consistently detected on western blots, both of which we interpreted as TAF15 (Figure 5; Figure S8; Figure S9). Although two bands were present in control samples—potentially reflecting post-translational modifications—the selective increase of the lower band following FUS+EWS double knockdown suggests that it may represent an alternatively spliced TAF15 isoform whose abundance is elevated upon concomitant FUS and EWS depletion. While this opens an interesting question for further investigation on the cross-regulation of the FET proteins, this was outside of the scope of our study, and thus, we focussed on the prominent TAF15 band at ∼77kDa. Five different anti-TAF15 antibodies raised in rabbit, ab133760 and ab13916 by Abcam, A300-307A and A300-308A by Bethyl and 28409 by Cell Signalling, demonstrated a single clear band whose intensity was decreased by TAF15 shRNA expression (Figure 5A-E, Figure S8A-E, Figure S9A-E; control vs TAF15 KD p<0.0001, control vs FUS+TAF15 KD p<0.0001, control vs EWS+TAF15 KD p<0.0001; control vs TAF15 KD p=0.0003, control vs FUS+TAF15 KD p=0.0006, control vs EWS+TAF15 KD p=0.0005; control vs TAF15 KD p=0.1812, control vs FUS+TAF15 KD p=0.3064, control vs EWS+TAF15 KD p=0.1753; control vs TAF15 KD p=0.0519, control vs FUS+TAF15 KD p=0.1071, control vs EWS+TAF15 KD p=0.0672; control vs TAF15 KD, p=0.0007, control vs FUS+TAF15 KD p=0.001, control vs EWS+TAF15 KD, p=0.0006).Neither of the Bethyl antibodies reached significance and A300-307A was the only antibody not to have the lower band, which could either be because the overall signal is too faint or because the epitope is not present in the putative alternatively spliced isoform. The only specific mouse anti-TAF15 antibody tested was MA3-078 by Invitrogen which presented the two TAF15 bands, with the main one intensity levels only decreasing when TAF15 expression was lowered with shRNA (Figure 5F, Figure S8F & Figure S9F: control vs TAF15 KD p<0.0001, control vs FUS+TAF15 KD p=0.0002, control vs EWS+TAF15 KD p<0.0001).

**Figure 5:**
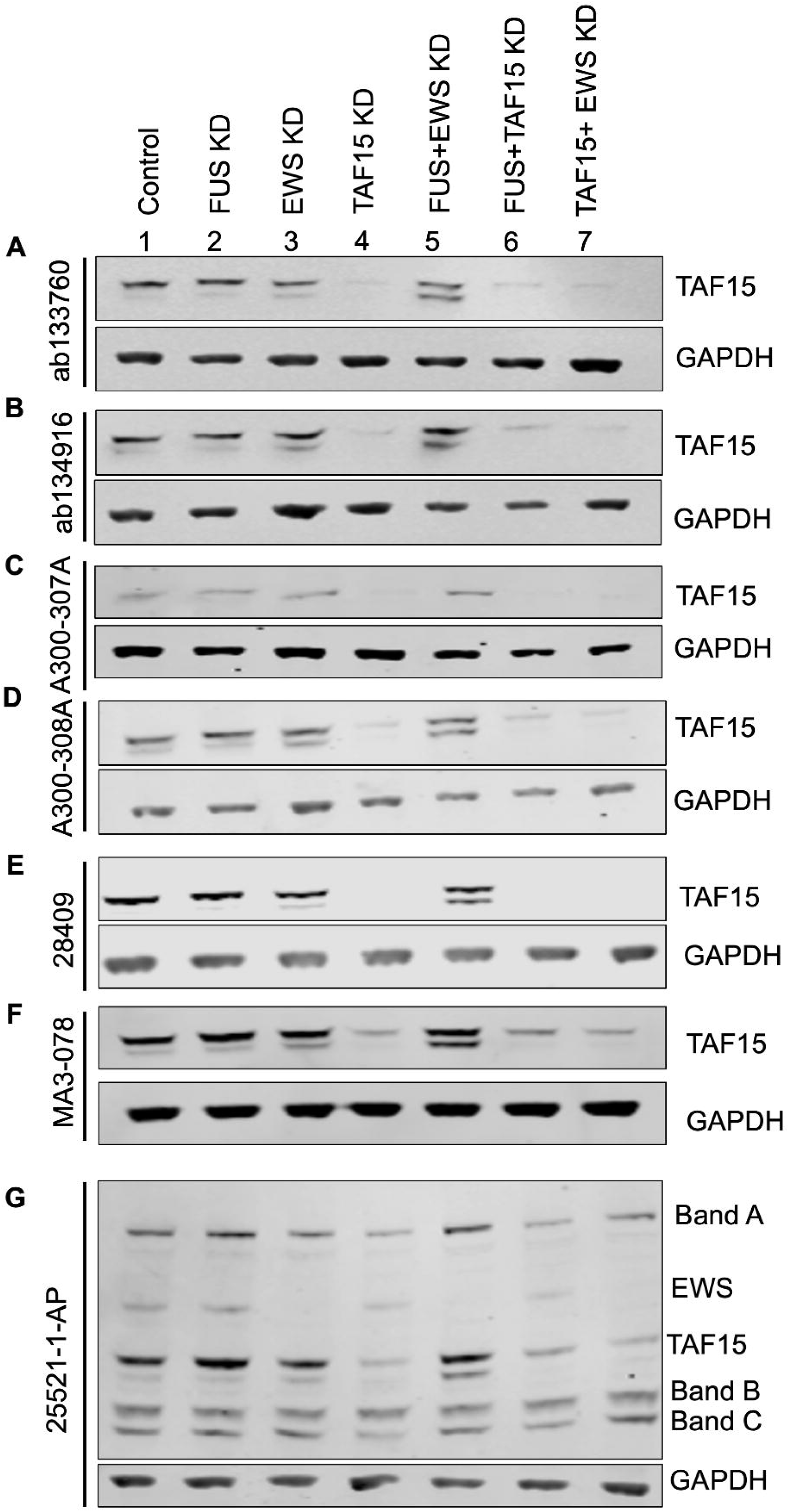
Anti-TAF15 antibody validation using single/double FET shRNA HeLa knockdowns. Western blot validation of seven commercially available anti-TAF15 antibodies showed a main band at ∼77 kDa corresponding to TAF15. Most antibodies (A, B, D, E, F & G) had an additional lower band determined to be TAF15. All antibodies show a decrease of the main TAF15 band in TAF15 only KD (Lane 4), FUS + TAF15 KD (Lane 6) and EWS + TAF15 KD (Lane 7). The lower band increased in the FUS+EWS KD (Lane 5). All bands were normalised to GAPDH loading control (37 kDa) and ratios of FET single/double knockdowns were then compared to control and significance determined by performing One-Way ANOVA with post-hoc Tukey’s multiple comparisons test (See Figure S8 for quantifications).

Finally, we tested two anti-TAF15 produced with multiple bands or none at all. Firstly, the rabbit antibody 25521-1- AP by ProteinTech, which presented with six different bands. Two of these bands were identified as being TAF15, at around 77kDa, with the main one significantly decreased in TAF15 KDs (Figure 5G, Figure S8G & Figure S9G; control vs TAF15 KD p<0.0001, control vs FUS+TAF15 KD p=0.0002, control vs EWS+TAF15 KD p=0.0001). Another one at around 90kDa when analysed was slightly altered when FUS levels were lowered and significantly decreased when EWS was knocked down (control vs FUS KD p=0.0395, control vs EWS KD p=0.0001, control vs FUS+EWS KD p=0.0003, control vs FUS+TAF15 p=0.0066, control vs EWS+TAF15 KD, p=0.0001). Three more bands (Band A, B & C, 120kDa, 60kDa and 55kDa respectively) showed no changes in any of the knockdowns suggesting that this antibody bound to the FET proteins as well as non-specific protein targets. The final anti-TAF15 antibody tested was sc-81121 by Santa Cruz which was tested at different dilutions, from 1:1000 to 1:100 but failed to show any band (Figure S9H).

Taken together, although the vast majority of tested antibodies performed well and showed substantial specificity for individual FET family members, a notable degree of antibody cross-reactivity with multiple FET proteins was still observed. While such cross-reactivity can be readily identified when proteins are resolved by molecular weight, it may lead to confounding results in applications where this separation is not possible, such as immunofluorescence (IF), immunohistochemistry (IHC) or immunoprecipitation (IP).

### FET antibodies show differing localisation patterns

To address the reported discrepancies in FET protein localisation, we employed only antibodies rigorously validated for specificity. Immunocytochemical analyses were conducted in two cellular models: rat primary neurons, representing a physiologically relevant neuronal system, and HeLa cells, widely utilised in FET protein research to date.

This dual approach facilitated cross-cell type and cross-species comparison to evaluate potential differences in staining patterns.

Overall, the anti-FUS antibodies show inconsistent nuclear/cytoplasmic sub-cellular detection between primary neurons and HeLa cells, but no drastic difference between antibodies targeting the N-terminal or C-terminal epitopes with all 7 anti-FUS antibodies showing both nuclear and cytoplasmic staining in primary cortical neurons, accompanied by poorer and more inconsistent staining in HeLa cells (Figure 6 with quantification in Figure S10). AMAb90549 and sc-373888 both show good nuclear signal with some cytoplasmic staining in primary neurons (mean nuclear intensity 89%) but they performed quite poorly in all replicates in HeLa cells. This was despite both working well in western blotting and having epitopes at opposite ends of the protein. Meanwhile BD Biosciences- 611384, and Atlas HPA008784, showed strong nuclear staining with some cytoplasmic staining in neurons that was recapitulated though to a lower extent in HeLa cells (both mean nuclear intensity 92%). However, whilst Novus NB100- 565 shows strong nuclear and some cytoplasmic staining in primary neurons (mean nuclear intensity 87%), in HeLa cells the nuclear staining is variable with some nuclei displaying no significant FUS signal. Proteintech 11570-1-AP and Novus NBP2-52874 show strong FUS nuclear staining in both neurons and HeLa cells accompanied by stronger cytoplasmic staining in both cell types (mean nuclear intensity 80% and 75%). There was no apparent difference in the level of cytoplasmic FUS seen when comparing the epitopes of the antibodies suggesting that this is not due to epitope availability.

**Figure 6:**
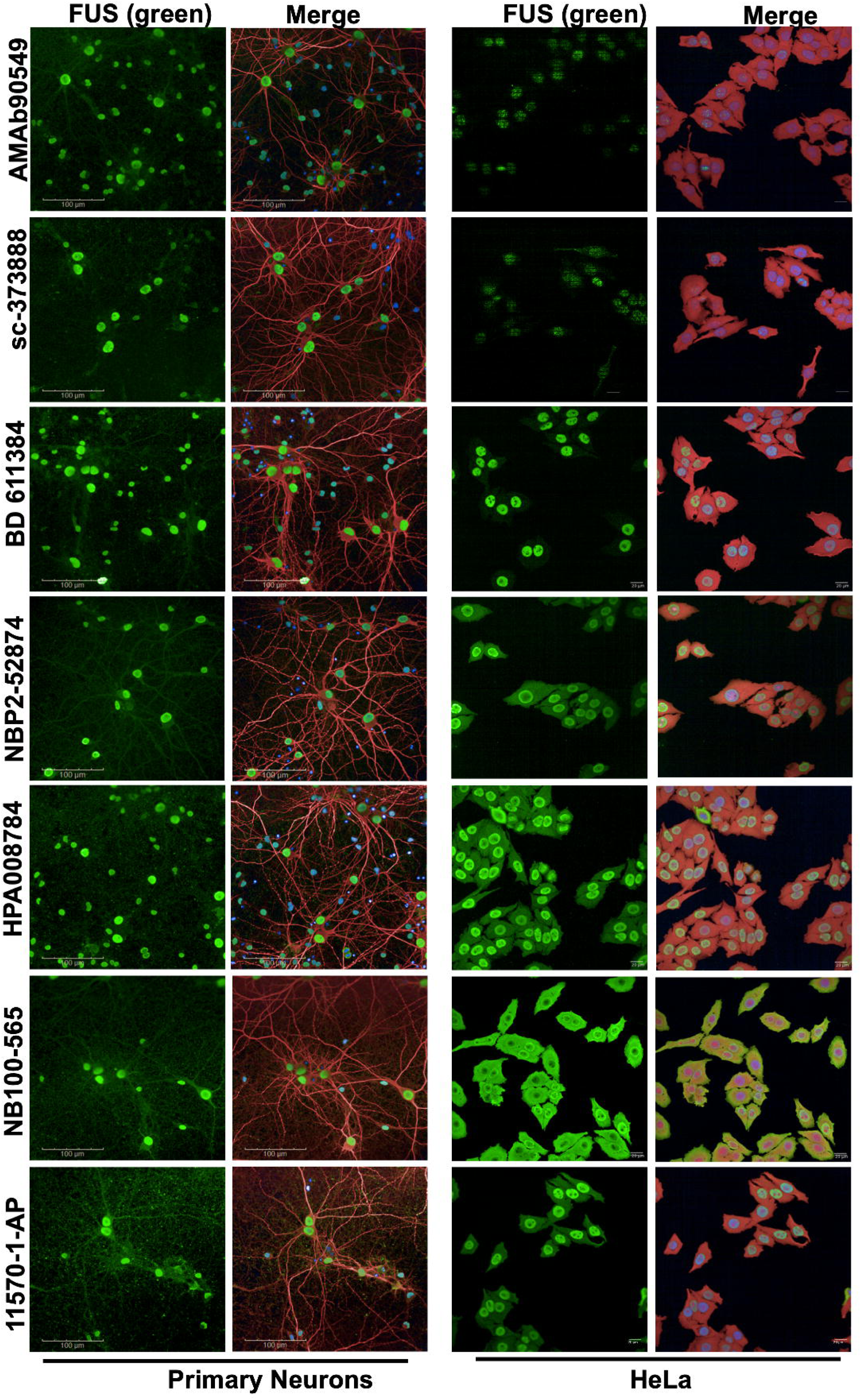
Immunocytochemistry validation of specific anti-FUS antibodies in primary cortical neurons and HeLa cells. anti-FUS antibody immunofluorescence detection of endogenous FUS (Green) in primary cortical neurons and wild-type HeLa cells. MAP2 (red) is used as a neuronal marker for primary cortical neurons, while Cell Mask (red) is used to stain the plasma membrane of HeLa cells. Nuclei are labelled with DAPI (blue). n = 3. 40x magnification. Scale bar (primary cortical neurons) = 100μm, Scale bar (HeLa cells)= 20μm. See Figure S10 for quantification graphs.

Out of the four anti-EWS antibodies, the AbCam ab133288, Santa Cruz sc-48404 and Santa Cruz sc-28327 performed well in immunocytochemistry experiments. In primary neurons, all three antibodies showed strong nuclear staining, which yielded the highest mean nuclear intensity of 94-95% compared to all FET antibodies tested (Figure 7, Figure S10 for quantification), which is similarly matched in HeLa Cells where EWS is almost entirely nuclear. Interestingly, in comparison the Proteintech 55191-1-AP failed to show the same strong nuclear localised staining with a clear diffuse cytoplasmic staining pattern reflected in a mean nuclear intensity of 48%. The western blot for this showed a series of very faint bands which may be due to a low level of non-specific binding that is more obvious in immunofluorescence. All epitopes for the EWS antibodies were in the N-terminus so not inference in terms of localisation and epitopes could be made.

**Figure 7:**
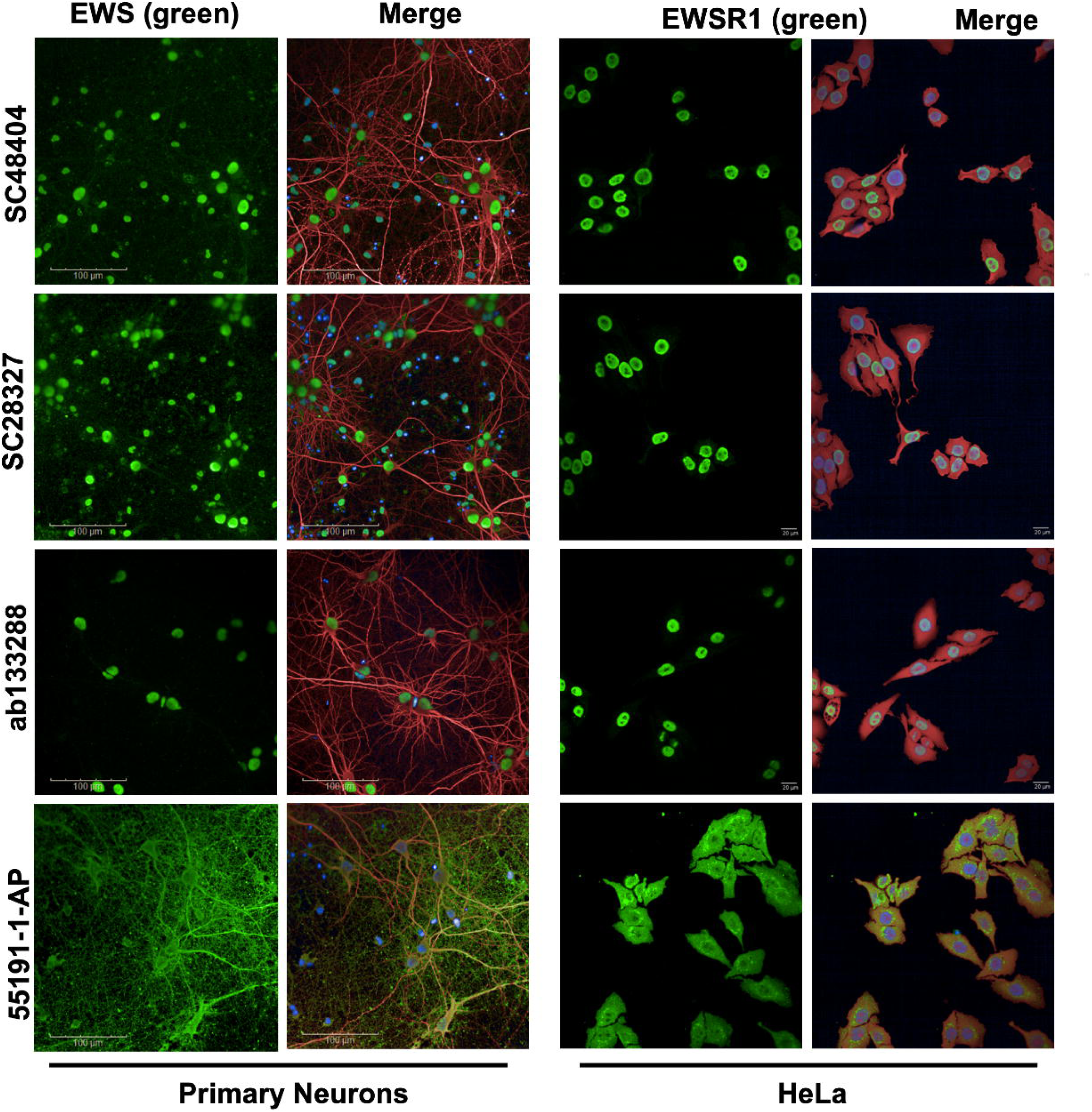
Immunocytochemistry validation of specific anti-EWS antibodies in primary cortical neurons and HeLa cells. anti-EWS antibody immunofluorescence detection of endogenous EWS (Green) in primary cortical neurons and wild-type HeLa cells. MAP2 (red) is used as a neuronal marker for primary cortical neurons, while Cell Mask (red) is used to stain the plasma membrane of HeLa cells. Nuclei are labelled with DAPI (blue). n = 3. 40x magnification. Scale bar (primary cortical neurons) = 100μm, Scale bar (HeLa cells)= 20μm. See Figure S10 for quantification graphs.

Anti-TAF15 antibodies showed generally consistent immunofluorescence staining in primary neurons, but variable staining in HeLa cells (Figure 8 and Figure S10 for quantification). The one antibody that did not perform well in primary neurons was AbCam ab133760, which in comparison to others showed much lower nuclear staining in neurons, which was reflected in the quantification (mean nuclear intensity 54%) though strong nuclear staining was observed in HeLa cells. This might be due to differences between species but as the epitope is unknown for this antibody it is not possible to confirm this. Both AbCam ab134916 and Bethyl A300-307A were similar with strong nuclear staining accompanied by some cytoplasmic staining in both cell types (mean nuclear intensity of 89%). The Bethyl A300-308A shows lower nuclear staining in neurons accompanied by higher cytoplasmic staining with a mean nuclear intensity of 74%. However, in HeLa cells, this is not recapitulated as there is no localised staining to the nucleus. Furthermore, the Cell Signalling-28409 anti-TAF15 antibody showed strong nuclear staining accompanied by some cytoplasmic staining with a mean nuclear intensity of 89%. Despite this, in HeLa cells, the same antibody was extremely inconsistent and sometimes failed to stain the nucleus alongside the high levels of staining in the cytoplasm. The final anti-TAF15 antibody was Invitrogen-MA3- 078 which showed strong nuclear staining in primary neurons accompanied by some cytoplasmic staining with a mean nuclear intensity of 86%. However, in HeLa cells, no nuclear staining was seen. Overall, these findings underscore the variability in antibody performance across cell types despite performing well in western blot, highlighting the importance of context-specific validation when selecting anti-TAF15 antibodies for experimental use.

**Figure 8:**
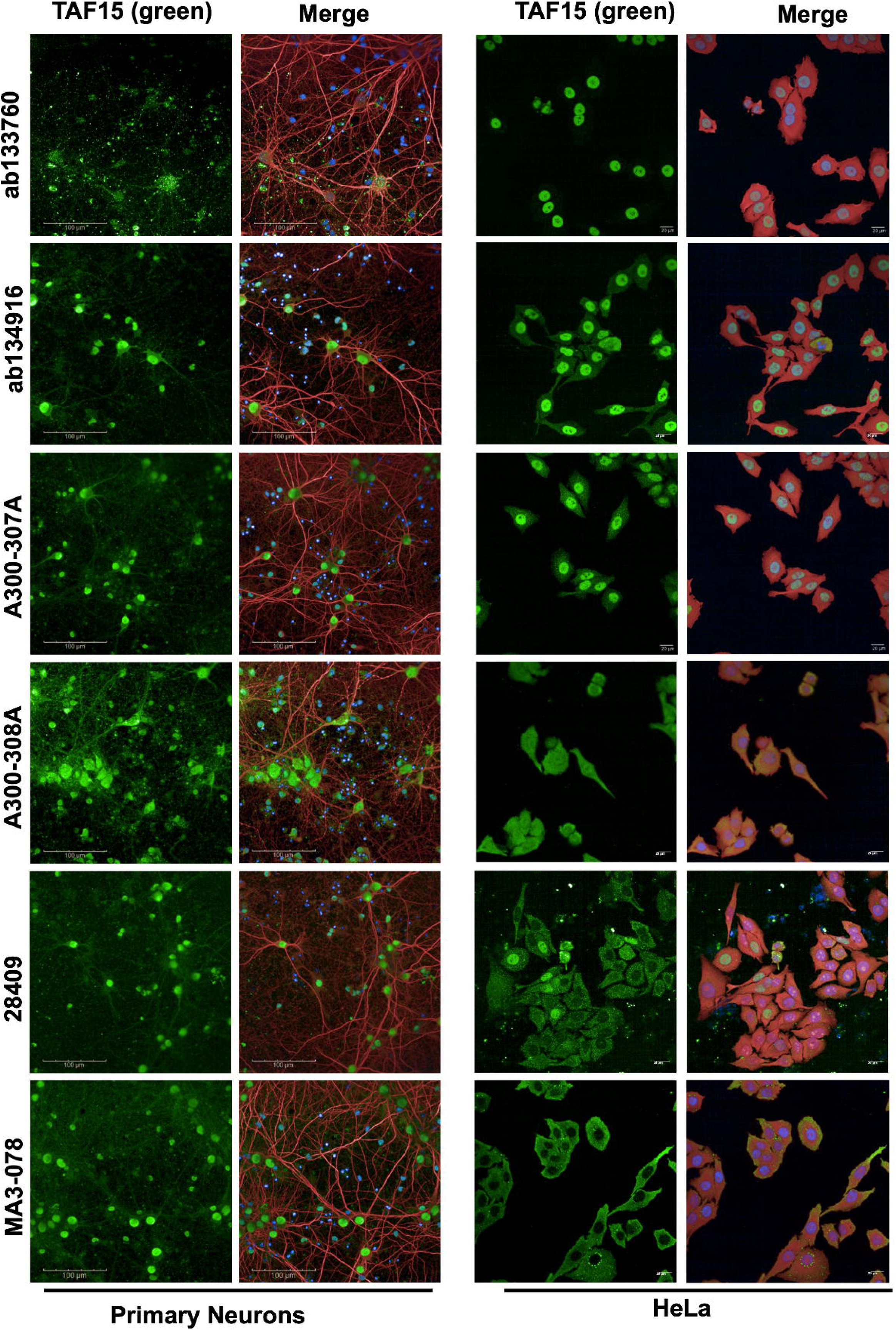
Immunocytochemistry validation of specific anti-TAF15 antibodies in primary cortical neurons and HeLa cells. anti-TAF15 antibody immunofluorescence detection of endogenous TAF15 (Green) in primary cortical neurons and wild-type HeLa cells. MAP2 (red) is used as a neuronal marker for primary cortical neurons, while Cell Mask (red) is used to stain the plasma membrane of HeLa cells. Nuclei are labelled with DAPI (blue). n = 3. 40x magnification. Scale bar (primary cortical neurons) = 100μm, Scale bar (HeLa cells)= 20μm. See Figure S10 for quantification graphs.

**Figure 9:**
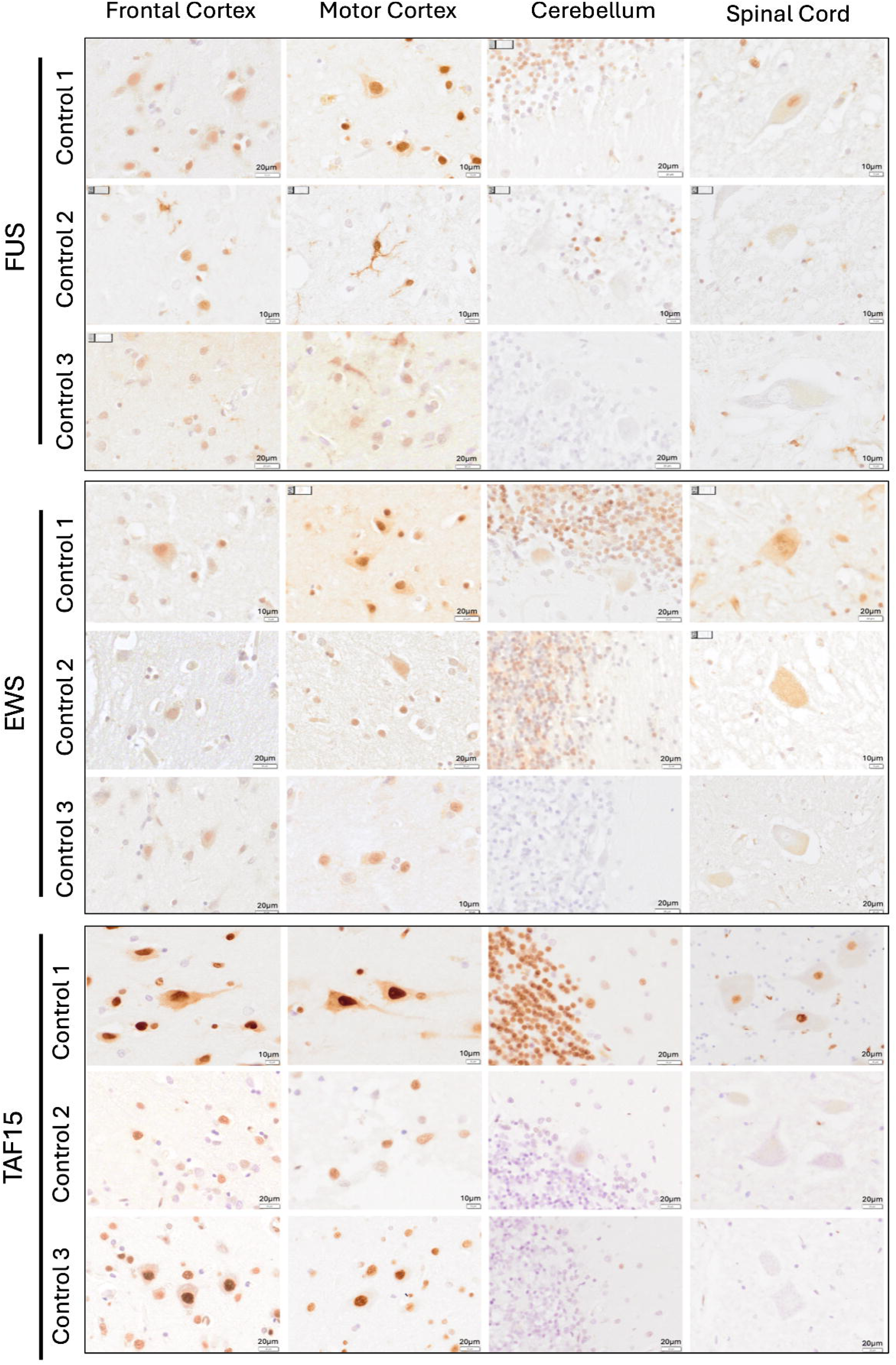
Immunohistochemistry-DAB of the FET proteins in postmortem tissue. Representative images of immunohistochemistry-DAB staining with anti-FUS Sigma-Aldrich-HPA008784, anti-EWS AbCam-ab133288 and anti-TAF15 AbCam-ab134916 (brown) in healthy human brain tissue. Nuclei are counterstained with Haematoxylin (blue). N=3, 40X magnification. Scale bars are indicated on each image (either 10μm or 20μm).

### FET protein distribution in healthy human tissue

Having assessed all of the antibodies by western blot and imaging, we chose one against each FET protein to assess their performance in human postmortem tissue to determine whether 1) the distribution of the FET proteins is similar across different brain regions and 2) how this distribution compares to the cell culture-based imaging results. Based on good precise selectivity and consistent imaging results across neurons and HeLa cells, we chose anti-FUS Sigma-Aldrich-HPA008784, anti-EWS AbCam-ab133288 anti-TAF15 AbCam-ab134916 antibodies. Immunohistochemistry was performed on sections from the frontal cortex, motor cortex, cerebellum and spinal cord from 3 different control donors (with no clinical evidence of any neurological condition and minimal/age-related pathology only). In all samples the FUS staining was strong in the nucleus with some cytoplasmic staining that was particularly increased in the motor cortex. This matches the strong nuclear with some cytoplasmic staining that was seen in primary neurons and HeLa cells. For EWS, there was clear nuclear staining in most samples with some cytoplasmic signal, albeit less pronounce compared to FUS. Again, this matches the cell-based imaging where EWS presents predominantly nuclear. For TAF15, there was a strong nuclear signal in most regions though this correlated with strong cytoplasmic staining. This was not consistent between samples but is in accordance with results from primary neurons and HeLa cells. Consistent across the antibodies, the staining does not show a strong signal in all regions of all samples. This might be due to expression levels which vary with age, or an artifact due to incomplete antigen retrieval during the tissue processing.

## Discussion

This study provides a comprehensive evaluation of antibody specificity within the FET protein family, emphasizing the risk of cross-reactivity and the necessity of rigorous antibody validation for reliable experimental outcomes. FUS, EWS, and TAF15 share high sequence homology and structural similarity, which complicates differentiation between family members. Our findings indicate that anti-FUS antibodies exhibit the greatest selectivity issues, with four out of eleven tested showing non-specific binding in single and double shRNA knockdown HeLa cells via western blotting. Notably, antibodies targeting epitopes at the extreme C-terminus of FUS demonstrated the highest cross-reactivity with EWS or both EWS and TAF15. One antibody (Proteintech-60160-1-Ig) detected all three FET proteins, suggesting that its epitope is highly conserved or identical across the family.

Selective anti-FUS antibodies produced strong nuclear staining with some cytoplasmic localisation in both cell types. Meanwhile all anti-EWS antibodies generally displayed good specificity by western blotting, though Proteintech-55191-1-AP showed faint unidentified background bands and performed poorly in immunocytochemistry, deviating from the predominantly nuclear staining patterns of other EWS-specific antibodies. Anti-TAF15 antibodies were largely selective by western blotting, though AbCam-ab133760, despite strong nuclear staining in HeLa cells, underperformed in neuronal immunocytochemistry. As a monoclonal antibody with an undisclosed epitope, species-specific sequence differences may explain this variability. Interestingly, two distinct bands were consistently observed for all anti-TAF15 antibodies, and quantification confirmed both correspond to TAF15. The increased expression of the second band following a FUS and EWS double knockdown suggests cross regulation occurs in the FET family. Its persistence suggests a previously uncharacterized isoform or post-translational modification with potential regulatory significance. Elucidating this phenomenon could uncover novel dimensions of FET protein biology, with implications extending to RNA metabolism and disease pathogenesis.

Across the FET family, in immunocytochemistry, no major differences were observed between N-terminal and C-terminal targeting antibodies; instead, staining variability appeared protein- and cell-type dependent. Differences between neurons and HeLa cells may reflect species-specific sequence variations or distinct cellular roles of FET proteins. Consistent with previous reports (Andersson et al., 2008), EWS was rarely detected in the cytoplasm compared to FUS and TAF15. Our findings align with a recent validation study (Alshalfie et al., 2023), which also identified Proteintech-60160-1-Ig as poorly selective despite its strong performance in immunoprecipitation. While our analysis spans western blotting, immunofluorescence, and immunohistochemistry, potential lot-to-lot variability remains an unaddressed limitation.

Detailed information on an antibody’s epitope is helpful in predicting potential cross-reactivity, allowing comparison with homologous regions in related proteins, yet it does not replace empirical validation and the diverging behaviour observed between HeLa cells and primary neurons underscores how strongly performance can depend on biological context. Establishing orthogonal benchmarks ultimately strengthens the translational relevance of findings and reduces misinterpretation in comparative studies.

Beyond validation, the functional consequences of antibody promiscuity demand careful scrutiny. While in this study, we did not assess antibodies in the context of Immunoprecipitation or other in vitro assays, cross-reactive antibodies can distort protein interaction networks or introduce off-target associations, introducing confounding variables that obscure true biological relationships. Addressing these risks will improve the integrity of mechanistic conclusions. Furthermore, reproducibility remains a pervasive challenge in research and particularly when it relies on the reliability of antibodies, underscoring the need for standardized quality control pipelines including inter-lot consistency testing. Implementing such frameworks across commercial suppliers would represent a transformative step toward harmonizing data quality in the field.

To our knowledge, this is the first systematic comparison of antibodies against all three FET proteins across knockdown models, providing a reference point that may help the field better understand antibody performance and potential sources of variability.

## Materials and Methods

### shRNA knockdowns

To deplete HeLa cells of FUS, EWS and TAF15, cells were seeded in 6-well plates at a density of 2.25x10^5^ cells/well in DMEM/F-12 with GlutaMAX™ supplement (Gibco, 10565018) supplemented with 10% Fetal Bovine Serum (Gibco, 26140079) and 1:100 Penicillin-Streptomycin (10,000 U/mL) (Gibco, 15140148) (Day0). The next day, 1 μg of shRNA plasmid per 6-well plate was transfected using TransIT-LT1 Transfection Reagent (Mirus, MIR2300) according to the manufacturer’s instructions. 24-hours after transfection (Day2), cells were split and replated using TrypLE (Gibco, 12605010) into a new 6-well plate, in a 1 to 2 ratio. The two following days (Day3 and Day4) medium was replaced with fresh media supplemented with 1.5 μg/mL Puromycin (Gibco, A1113803).

### Cell Lysates

On Day 5 post transfection, selection was stopped. After ∼6 hours, cells were harvested. by trypsinisation with TrypLE (GibcoTM, 12605010) followed by washing with DPBS and harvest of cell pellets by centrifugation at 500 x g for 5 minutes. Cells were resuspended in 100 μL/well RIPA buffer supplemented with cOmplete™ Protease Inhibitor Cocktail (Roche, 4693116001), followed by incubation on ice for 20 minutes and centrifugation at 16,000 x g for 20 minutes at 4°C to remove cell debris. Supernatant was transferred to new 1.5 mL Eppendorf tube, mixed 1:1 with 2x sample buffer (4% SDS, 20% glycerol, 120mM Tris pH6.8) and boiled at 95°C for 5 minutes.

### qPCR

On Day 5 post transfection, the HeLa the cells cells were harvested by trypsinisation with TrypLE (Gibco, 12605010) followed by washing with DPBS and harvest of cell pellets by centrifugation at 500 x g for 5 minutes. Pellets were resuspended in 300 μl (for up to 106 cells) of TRISure (Bioline Reagents Ltd., BIO-38032) supplemented with 10 μl β-mercaptoethanol (ITW Reagents, A1108) per ml TRIsure. RNA was isolated using the Direct-zolTM RNA Miniprep Plus Kit (ZYMO RESEARCH, R2072) according to the manufacturer’s instructions and eluted in 50 μl of RNase-free water. Reverse Transcription of up to 2 μg of RNA into cDNA was performed using the LunaScript RTSuperMix Kit (New England Biolabs, E3010). SYBR Green RT-qPCR reactions were carried out to validate the Single/Double FET Knockdowns in HeLa cells using the PowerUp SYBR Green Master Mix (Applied Biosystems, A25742). Pipetting was completed by the QIAgility HEPA/UV automated pipetting robot, with 32ng cDNA and a final concentration of primers of 600nM per 20 μl reaction. Both PCR amplification and fluorescence detection were conducted using the Rotor-Gene Q thermocycler. Melting curve analyses were conducted to confirm and assess the specificity of each primer used.

### Western blotting

Samples were run on an 10% NuPAGE Bis-Tris Polyacrylamide Midi Gel (Thermo Fisher, WG1203BOX) along with a Precision Plus Protein Dual Color Standards ladder (Bio-Rad, 1610374) using 1x MOPS SDS Running Buffer (Invitrogen, NP0001). The gel was removed from the casting apparatus and equilibrated in 50 ml of transfer solution consisting of 2× transfer buffer (Invitrogen, NP00061) and 5 ml of methanol, brought to a final volume of 50 ml with distilled water. Dry transfer onto a nitrocellulose membrane (Invitrogen, IB23001) was conducted using the iBot2 Gel transfer device (Invitrogen, IB21001) according to the manufacturer’s instructions. The membrane was blocked in 5% non-fat dried milk in TBST (500mM Trizma® base (Sigma-Aldrich, T4661), 1.54mM NaCl (Sigma-Aldrich, S5886), pH 7.6, 0.1% Tween20 (Sigma Aldrich P9416)) for 30 minutes at room temperature, followed by incubation with the respective FET primary antibodies in 1% Milk TBST overnight at 4°C with mouse anti-GAPDH (1:5000 Sigma-Aldrich, G8795). After washing the blot for 3 x 10 minutes with TBST secondary Dylight antibodies (1:5,000, Anti-Rabbit Goat DyLight 800 (Thermo Fisher SA5-35571), Anti-Mouse Goat DyLight 680 (Thermo Fisher (35518)) in 1% Milk TBST were added for 1 hour, followed by TBST washes. Membranes were scanned using the Odyssey CLx Imaging System (LI-COR) and analysed using Image Studio Lite (LI-COR).

### Primary neurons culture

The day before plating, 96-well plates were coated with ∽100 μL of 0.1 mg/mL Poly-D-Lysine hydrobromide (Sigma-Aldrich, P6407) and kept at 37°C overnight before washing with sterile ddH2O and left to dry at RT until needed. Primary cortical neurons were obtained from E18 rat pups. Briefly, cortices were dissected into HBSS (-calcium and magnesium, Thermo Fisher, 14170112) with 2.5% Trypsin, left to incubate at 37°C for 30 minutes followed by DNaseI treatment (Thermo Fisher, 18047019) in HBSS with calcium and magnesium (Thermo Fisher, 24020117). The supernatant was aspirated, removing the HBSS, trypsin and DNaseI. The cells were resuspended in triturating solution with Neurobasal Complete medium then put through a 70 μM cell strainer and counted using a hemacytometer, 10 μL of the neurons was added to 90 μL of Trypan Blue (Thermo Fisher, 15250061). Neurons were plated at a confluency of 18,000 cells/well in a 96-well plate. Neurons were maintained in culture in Neurobasal Complete medium in an incubator at 37°C and 5% CO2 until fixation.

### Immunocytochemistry

On DIV21, rat primary neurons were fixed in 100 μL/well of 4% Paraformaldehyde (PFA) for 10 minutes, followed by ice-cold methanol for 10 minutes. HeLa cells were maintained as for the knockdown experiments for 5 DIV before fixing in 100 μL/well 4% PFA for 10 minutes. The same protocol was used for both cell types. Fixed cells were firstly washed three times in DPBS (Gibco, 14040133), permeabilised and unspecific antibody binding blocked, by incubation in 1% goat serum in 0.2% Triton X – PBS for 30 minutes at RT. Primary antibodies against the specific FET proteins and chicken MAP2 (for the primary neurons, Antibodies.com A85363) were then diluted in goat serum with 0.2% Triton X – PBS and incubated overnight at 4°C. The following day, cells were washed three times for 5 minutes in DPBS and incubated with the relevant secondary antibodies (Invitrogen anti-rabbit 488 (A11008), anti-mouse 488 (A11029) and anti-chicken 633 (A21103)) diluted in 2% goat serum with 0.2% Triton X – PBS for 1 hour at RT. Cells were washed again three times in DPBS and a drop of DAPI (Thermo Fisher, 62248) was then added to each well or coverslip. HeLa cells were additionally stained with CellMask Deep Red (1:10000, Thermo Fisher H32721). Cells were washed once more in DPBS before imaging.

### Immunohistochemistry

Post-mortem human tissue was provided by the London Neurodegenerative Diseases Brain Bank, King’s College London. Sections of 7μm thickness were cut from formalin-fixed paraffin-embedded tissue blocks, and subject to standard immunohistochemistry protocol. Citrate buffer microwave pre-treatment was conducted to enhance antigen retrieval. After blocking in 2.5% normal horse serum, sections were incubated with primary antibody diluted in Animal-Free blocker (Vector Laboratories, Cat.SP5035) at room temperature for one hour. Antibodies used: FUS - HPA008784 (Sigma-Aldrich), EWS 133288 (AbCam), TAF 15 134916 (AbCam) all at 1:100. Following washes, sections were incubated with biotinylated secondary antibody in ImmPRESS HRP Horse Anti-Rabbit IgG Polymer, Peroxidase (Vector Laboratories, Cat.MP-7401) for 30 minutes. The sections were then washed and incubated for 3-5 minutes in 3,3′-diaminobenzidine chromogen, with the solution prepared according to manufacturer’s instructions (Vector Laboratories, Cat.SK-4103). Finally, sections were counterstained with Harris’ haematoxylin, dehydrated in IMS and Xylene and coverslips mounted.

### Imaging

Both Primary neurons and HeLa cells were imaged on the Opera Phenix (Perkin Elmer) microscope (Wohl Cellular Imaging Centre). The 96-well plates were scanned at 40x magnification using the water objective lens. 10 Z-stacks of 0.2 μm steps apart were obtained for the DAPI, green (488 nm) and far-red (633 nm) channels. Once scanned, the Harmony software was used to create 2D analysis pipelines to carry out quantitative comparisons of sub-cellular localisation between different FET antibodies in Primary neurons (see figure S11 for examples). DAB Human slides were imaged using an Olympus Slide Scanner VS200. Detailed single-plane images were acquired at 40x magnification with a High Focus Position Density, including prefocus on both sample and non-sample regions to allow greater accuracy.

### Plasmids

pSUPuro plasmids were cloned by inserting double-stranded oligos into pSUPuro between the BglII and HindIII sites as described in (Brummelkamp et al., 2002; Paillusson et al., 2005). The shRNAs expressed from pSUPuro FUS t4 and t5 are targeting nucleotides 1143–1161 (GTAAAGAATTCTCCGGAAA) and nucleotides 614-632 (GATCAATCCTCCATGAGTA) of FUS mRNA, numbering according to NM_004960. The shRNAs expressed from pSUPuro EWS t1 and t2 are targeting nucleotides 828–826 (CAGCCAAGCTCCAAGTCAA) and nucleotides 717-735 (GCAGAACACCTATGGGCAA) of EWS mRNA, numbering according to NM_013986. The shRNAs expressed from pSUPuro TAF-15 t1 and t2 are targeting nucleotides 1822–1840 (GAAGAAACGACTACAGAAA) and nucleotides 1481-1499 of TAF15 mRNA, numbering according to NM_139215. The shRNA expressed from pSUPuro-IRES, which served as negative control targets the sequence TGTGTTTAGTCGAGGTTAA, present in the Encephalomyocarditis virus (EMCV) Internal Ribosome Entry Site (IRES).

### Oligonucleotides

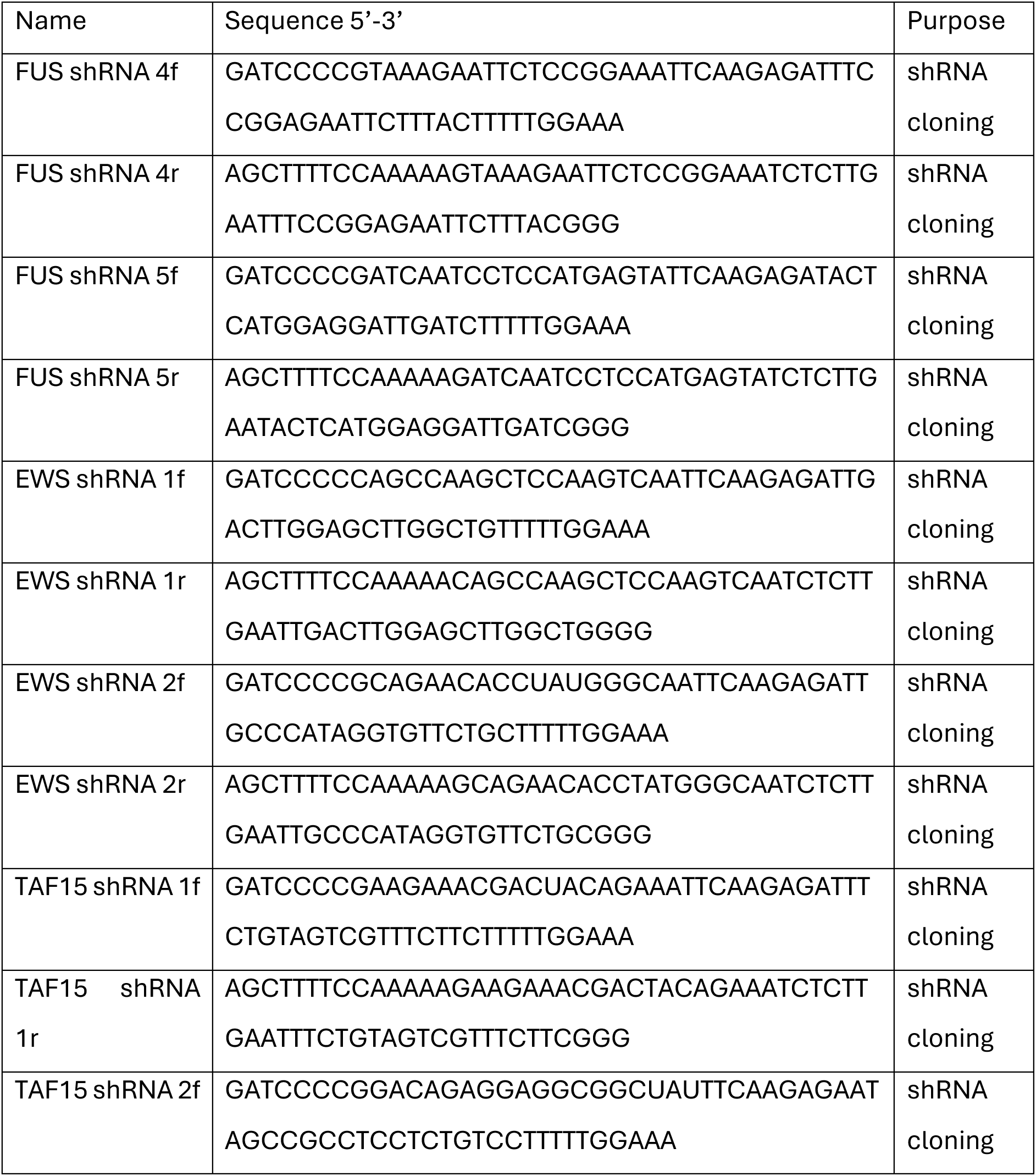
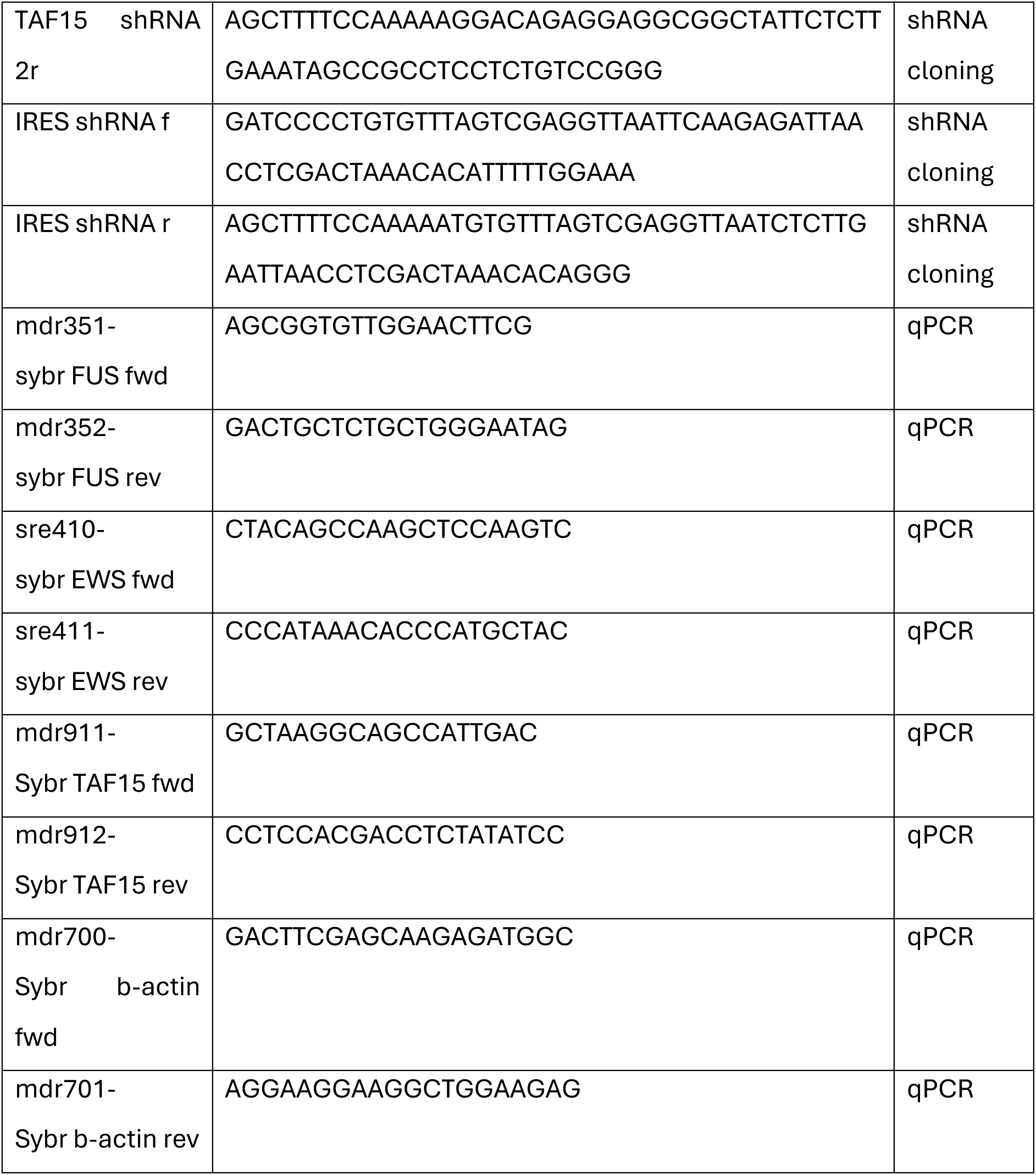

### Human Post-mortem tissue samples

| BBN-ID | Label | Age | Sex | PMD (hours) | Fixation time<br>(weeks) | Diagnosis |
| --- | --- | --- | --- | --- | --- | --- |
| BBN002.30136 | 1 | 90 | F | 48 | 5 | control |
| BBN002.36907 | 2 | 99 | F | 45.5 | 10 | Control |
| BBN 13802 | 3 | 74 | M | 22.5 | 14 | control |

## Acknowledgements

The authors would like to thank Dr George Chenell at the Wohl Cellular Imaging Centre (Institute of Psychiatry, Psychology and Neuroscience, King’s College London) for their help with training and image acquisition.

