## Supplementary Data for "A validated panel of commercial antibodies for reliable detection of FET proteins"

### Supplementary Material

Tacconelli et al, 2025

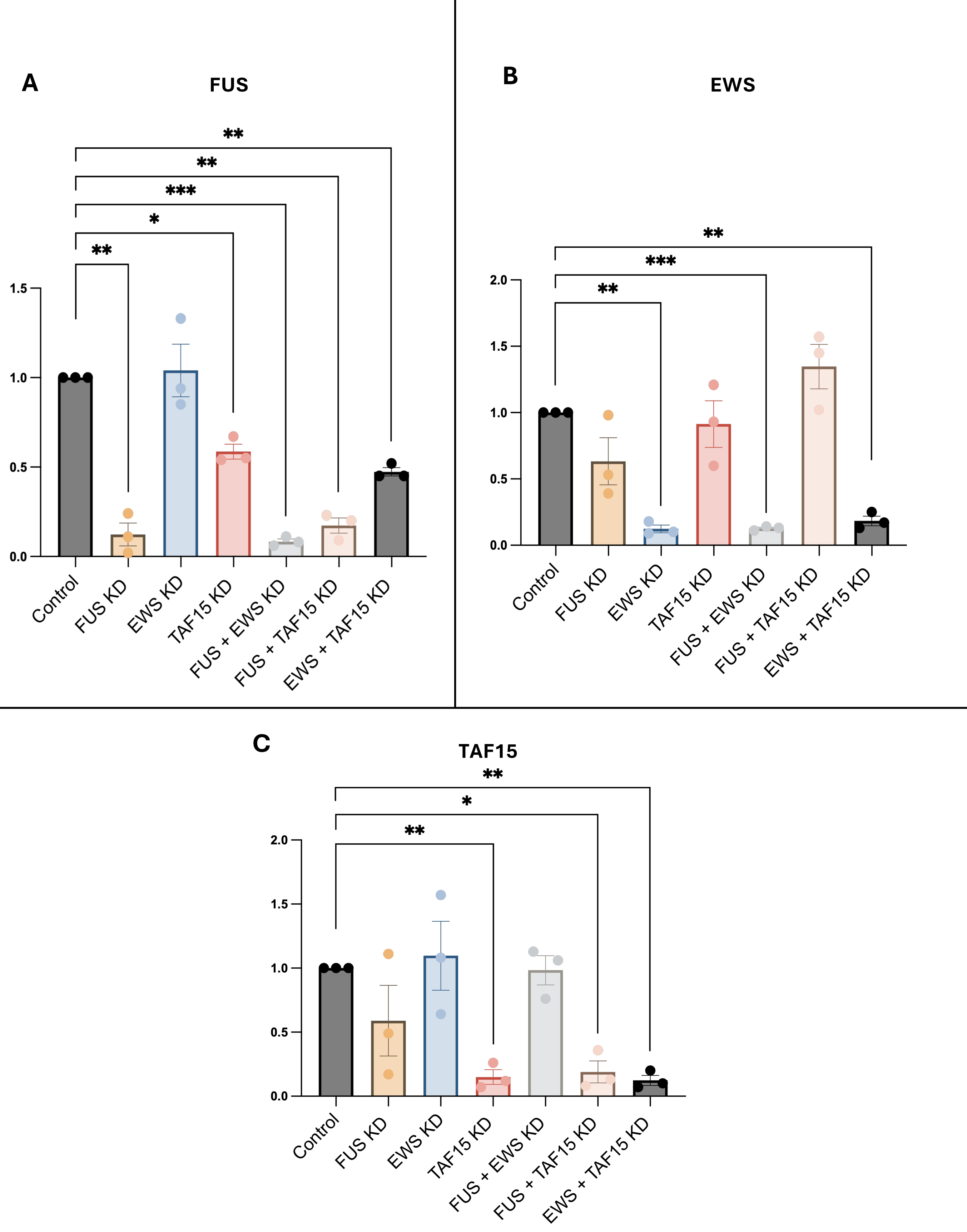


**Figure S1: RT-qPCR validation of single/double FET shRNA knockdown in HeLa cells.**

FUS, EWS and TAF15 mRNA levels in single and double FET shRNA knockdown in HeLa cells are shown. A) FUS mRNA levels B) EWS mRNA levels and C) TAF15 mRNA levels. HeLa cells were harvested for each condition and RNA isolated from total lysates for RT-qPCR. Ct values were normalised to control and Actin. *($p \leq 0.05$), **($p \leq 0.01$), ***($p \leq 0.001$). Mean ± SEM is plotted. n = 3.

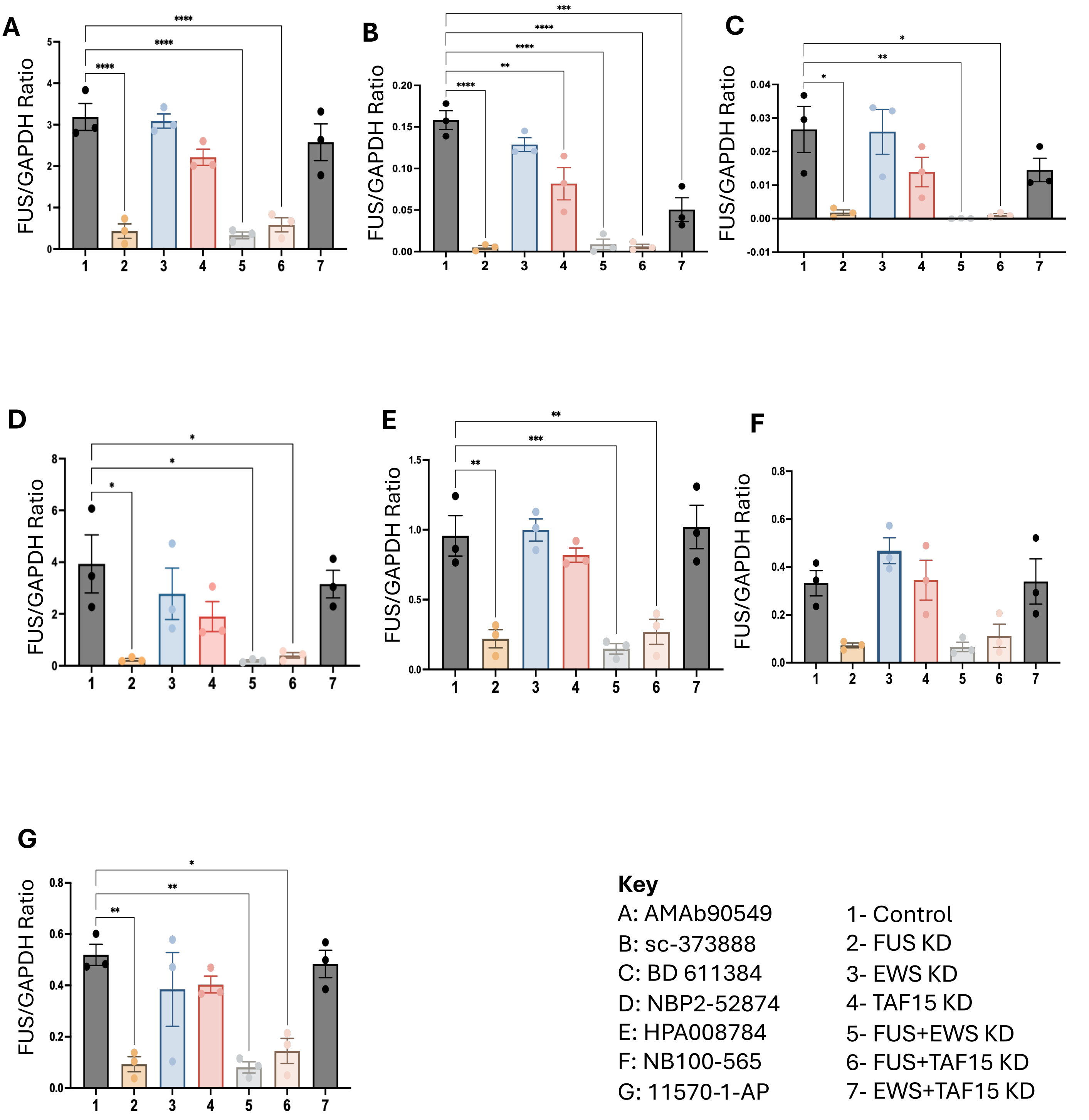


**Figure S2: Quantification of western blots for specific anti-FUS antibodies.** Western blot of seven commercially available anti-FUS antibodies showed a single band at ~70 kDa corresponding to FUS (Figure 2). All antibodies show a decrease of FUS protein levels in FUS only KD (Lane 2), FUS + EWS KD (Lane 5) and FUS + TAF15 KD (Lane 6). All bands were normalised to GAPDH loading control (37 kDa) and ratios of FET single/double knockdowns were then compared to control and significance determined by performing One-Way ANOVA with post-hoc Tukey's multiple comparisons test. *($p \leq 0.05$), **($p \leq 0.01$), ***($p \leq 0.001$), ****($p \leq 0.0001$). Mean ± SEM is plotted. n=3.

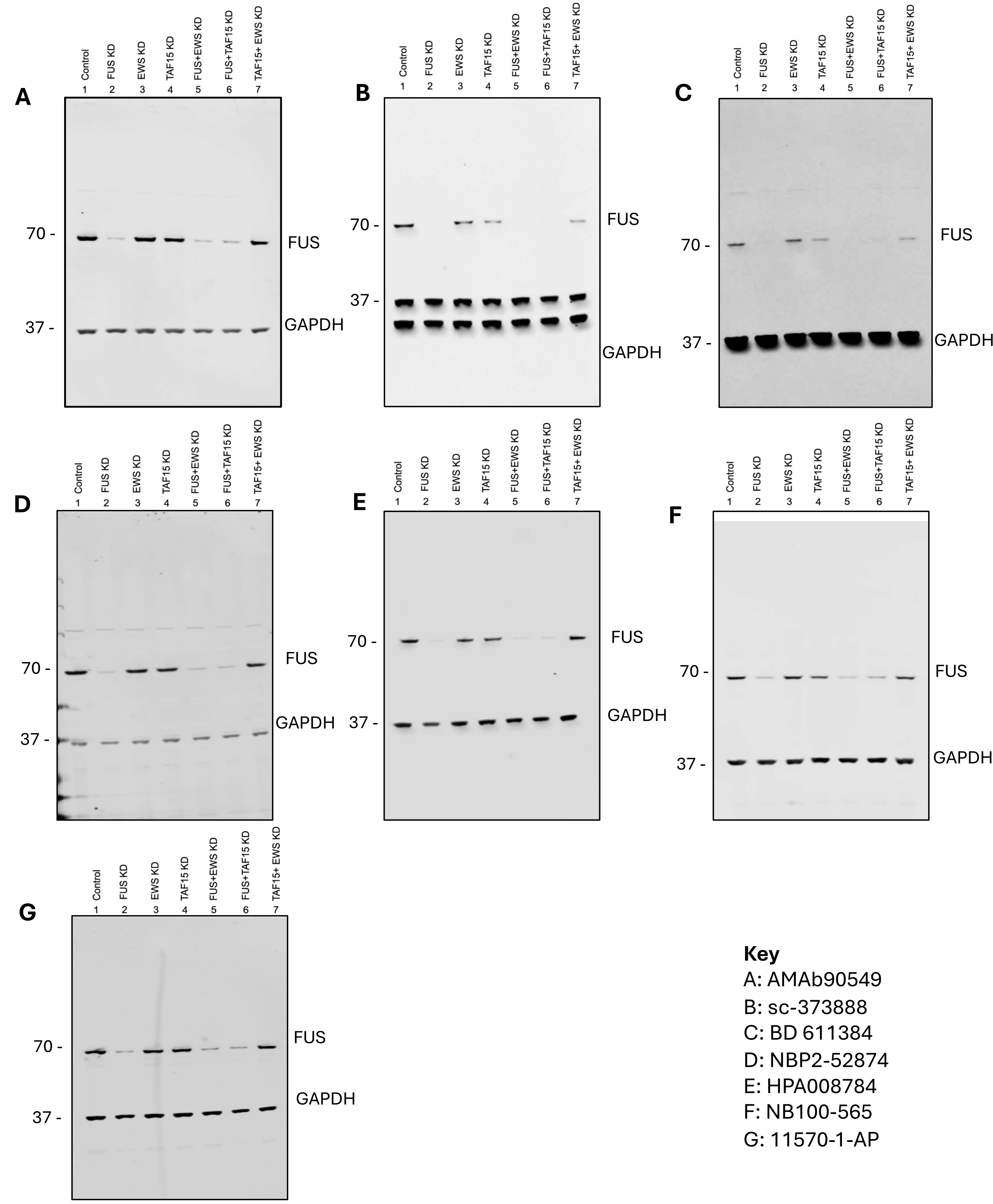


**Figure S3: Specific anti-FUS antibodies confirmed by single/double FET shRNA HeLa knockdowns.** Full Western blot validation of seven commercially available anti-FUS antibodies showed a single band at ~70 kDa corresponding to FUS. All antibodies show a decrease of FUS protein levels in FUS only KD (Lane 2), FUS + EWS KD (Lane 5) and FUS + TAF15 KD (Lane 6). All bands were normalised to GAPDH loading control (37 kDa) and ratios of FET single/double knockdowns were then compared to control and significance determined by performing One-Way ANOVA with post-hoc Tukey's multiple comparisons test. Please note for B a different lot of the GAPDH antibody was used which resulted in a double band. (see Figure S2 for quantification).

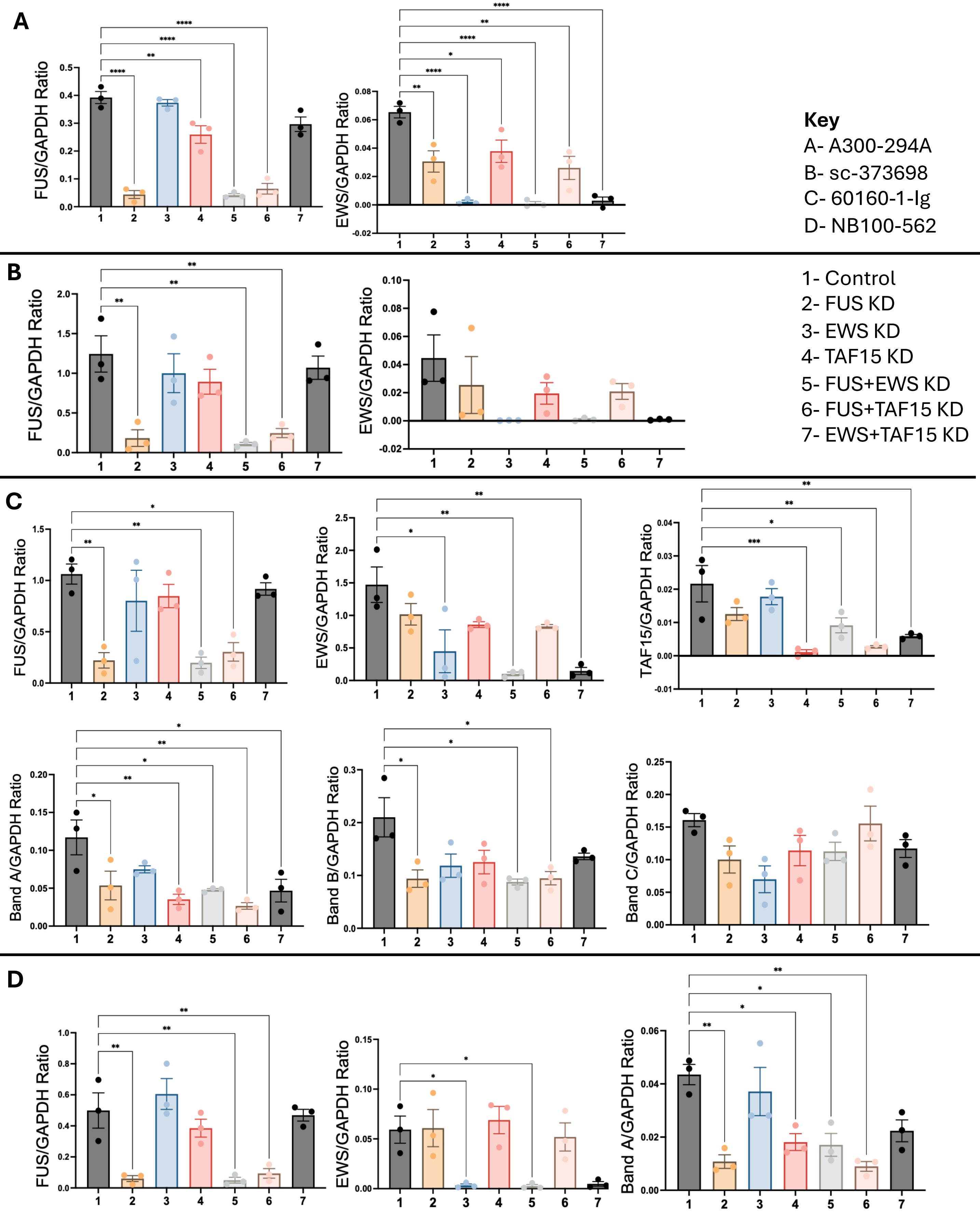


**Figure S4: Quantification of western blots for non-specific anti-FUS antibodies.** All antibodies show a significant decrease of FUS protein levels in FUS only KD (Lane 2), FUS + EWS KD (Lane 5) and FUS+TAF15 KD (Lane 6). However, all four showed additional bands identified as EWS and/or TAF15 as well as as yet unidentified bands. All bands were normalised to GAPDH loading control (37 kDa) and ratios of FET single/double knockdowns were then compared to control and significance determined by performing One-Way ANOVA with post-hoc Tukey's multiple comparisons test. *(p ≤0.05), **(p ≤0.01), ***(p ≤0.001), ****(p ≤0.0001). Mean ± SEM is plotted. n=3.

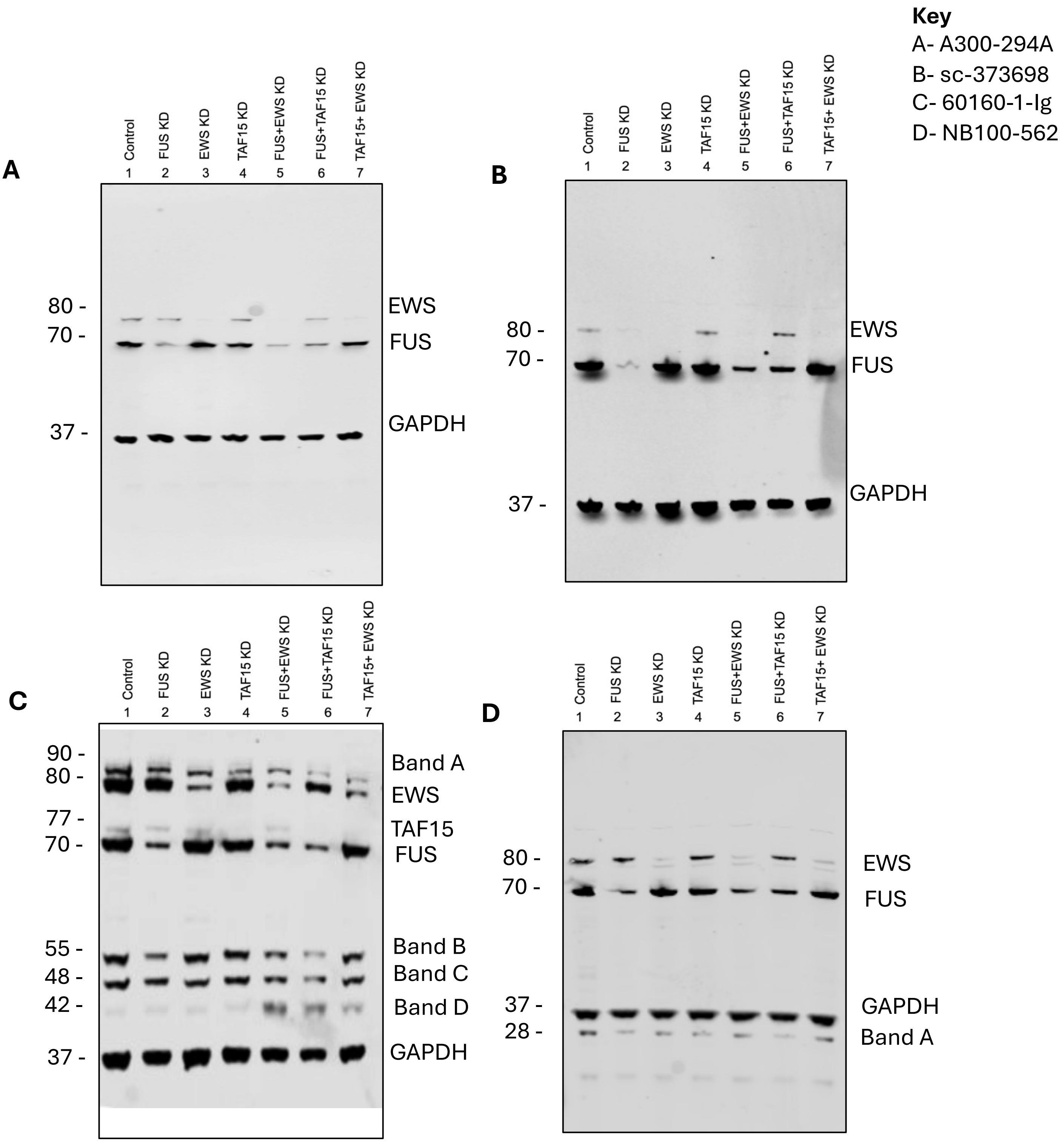


**Figure S5: Cross-reactivity of some anti-FUS antibodies confirmed by single/double FET shRNA HeLa knockdowns.** Full Western blot validation of four commercially available anti-FUS antibodies showed a single band at ~70 kDa corresponding to FUS. All antibodies show a significant decrease of FUS protein levels in FUS only KD (Lane 2), FUS + EWS KD (Lane 5) and FUS+TAF15 KD (Lane 6). However, all four showed additional bands identified as EWS and/or TAF15 as well as additional unidentified bands. All bands were normalised to GAPDH loading control (37 kDa) and ratios of FET single/double knockdowns were then compared to control and significance determined by performing One-Way ANOVA with post-hoc Tukey's multiple comparisons test (See Figure S4 for quantifications).

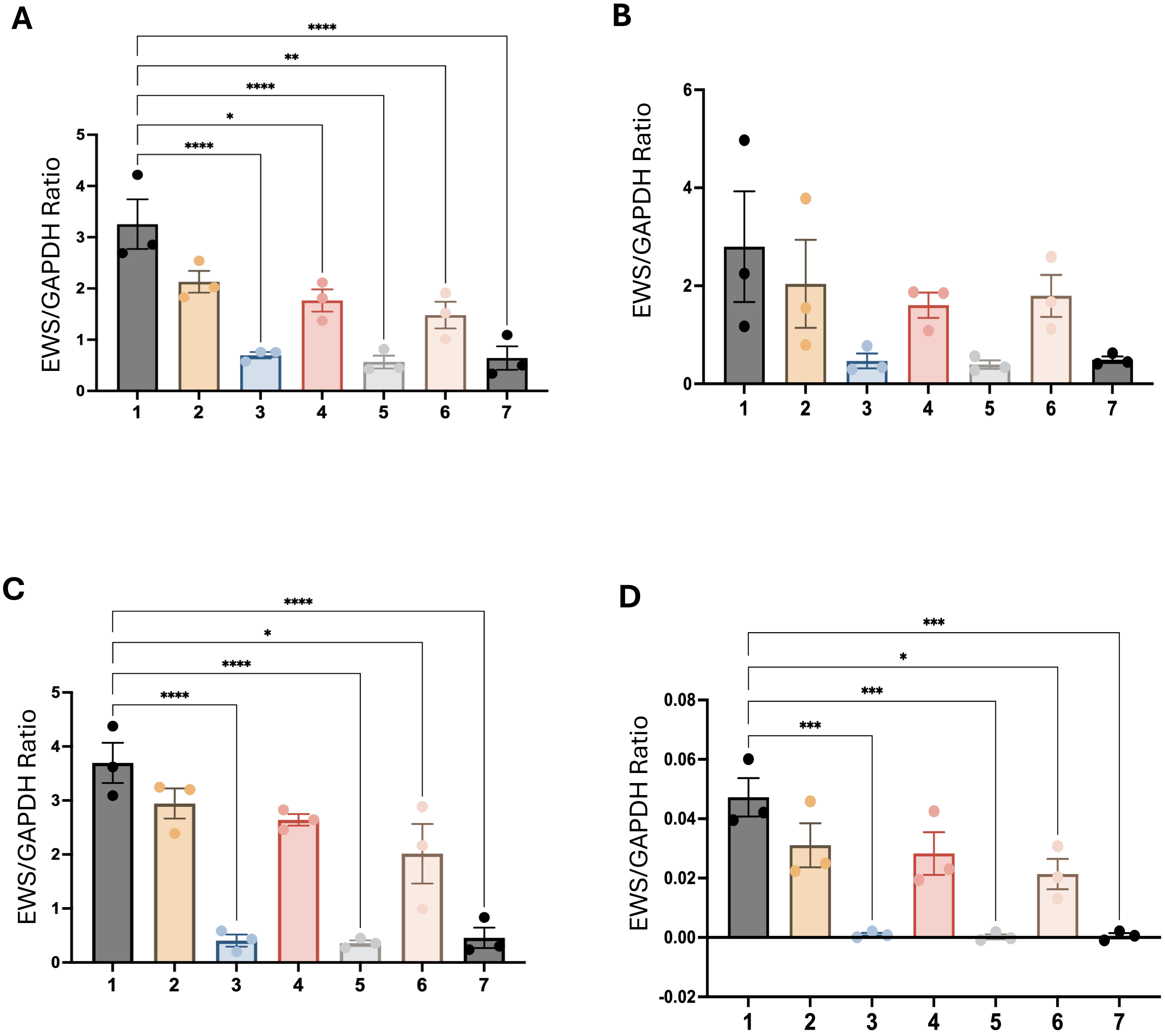


**Key**
A: sc48404
B: sc28327
C: ab133288
D: 55191-1-AP

1- Control
2- FUS KD
3- EWS KD
4- TAF15 KD
5- FUS+EWS KD
6- FUS+TAF15 KD
7- EWS+TAF15 KD

**Figure S6: Quantification of western blots for anti-EWS antibodies.** All antibodies show a decrease of EWS protein levels in EWS only KD (Lane 3), FUS + EWS KD (Lane 5) and TAF15 +EWS KD (Lane 7). All bands were normalised to GAPDH loading control (37 kDa) and ratios of FET single/double knockdowns were then compared to control and significance determined by performing One-Way ANOVA with post-hoc Tukey's multiple comparisons test (See Figure S6 for quantifications). *($p \leq 0.05$), **($p \leq 0.01$), ***($p \leq 0.001$), ****($p \leq 0.0001$). Mean ± SEM is plotted. n=3.

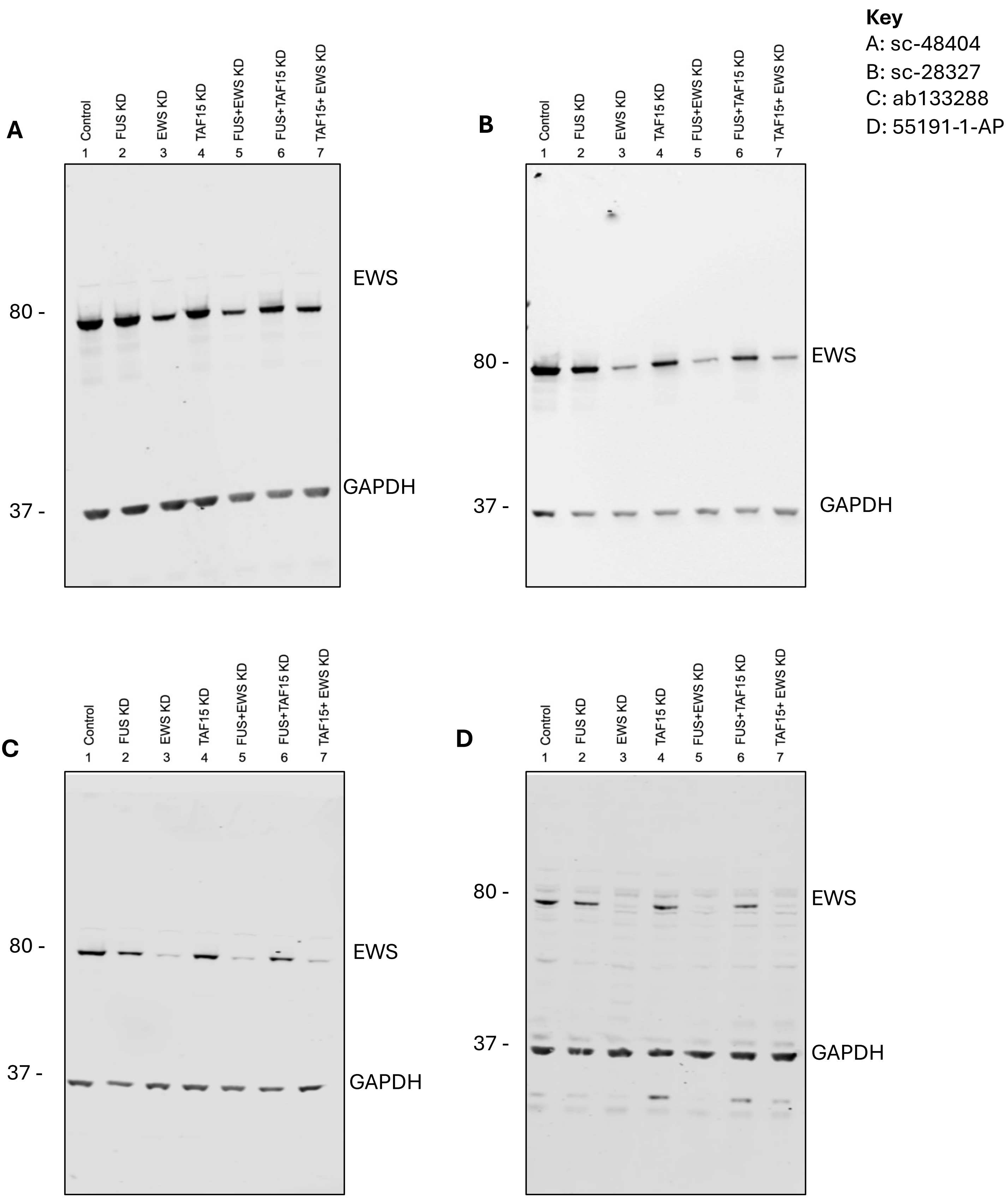


**Figure S7: Specific anti-EWS antibodies confirmed by single/double FET shRNA HeLa knockdowns.** Full Western blot validation of seven commercially available anti-EWS antibodies showed a single band at ~80 kDa corresponding to EWS. All antibodies show a significant decrease of EWS protein levels in EWS only KD (Lane 3), FUS + EWS KD (Lane 5) and TAF15 +EWS KD (Lane 7). All bands were normalised to GAPDH loading control (37 kDa) and ratios of FET single/double knockdowns were then compared to control and significance determined by performing One-Way ANOVA with post-hoc Tukey's multiple comparisons test (See Figure S6 for quantifications).

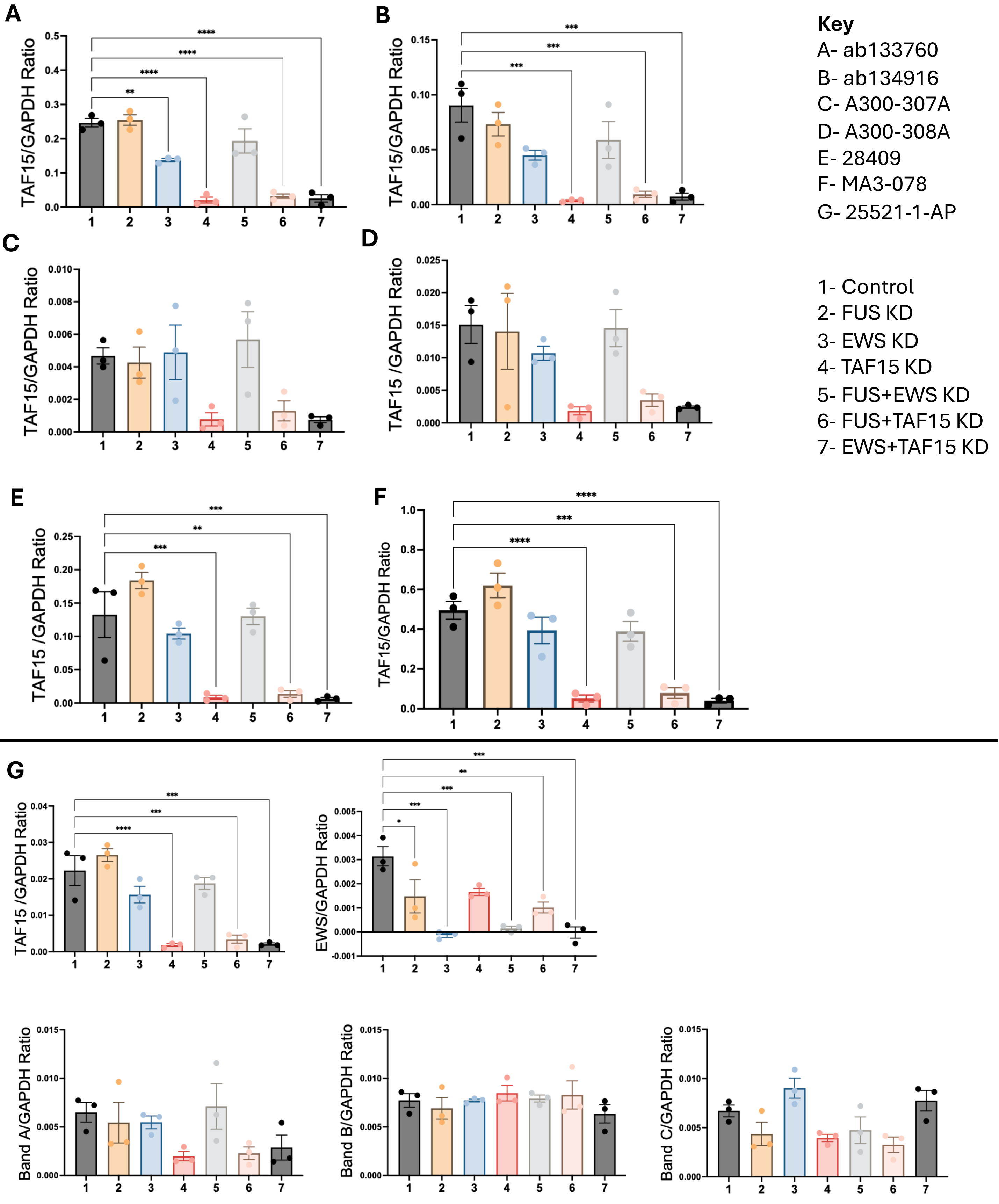


**Figure S8: Quantification of anti-TAF15 blots .** All antibodies show a decrease of the main TAF15 band in TAF15 only KD (Lane 4), FUS + TAF15 KD (Lane 6) and EWS + TAF15 KD (Lane 7). The lower band increased in the FUS+EWS KD (Lane 5). All bands were normalised to GAPDH loading control (37 kDa) and ratios of FET single/double knockdowns were then compared to control and significance determined by performing One-Way ANOVA with post-hoc Tukey's multiple comparisons test. ). *(p ≤0.05), **(p ≤0.01), ***(p ≤0.001), ****(p ≤0.0001). Mean ± SEM is plotted. n=3.

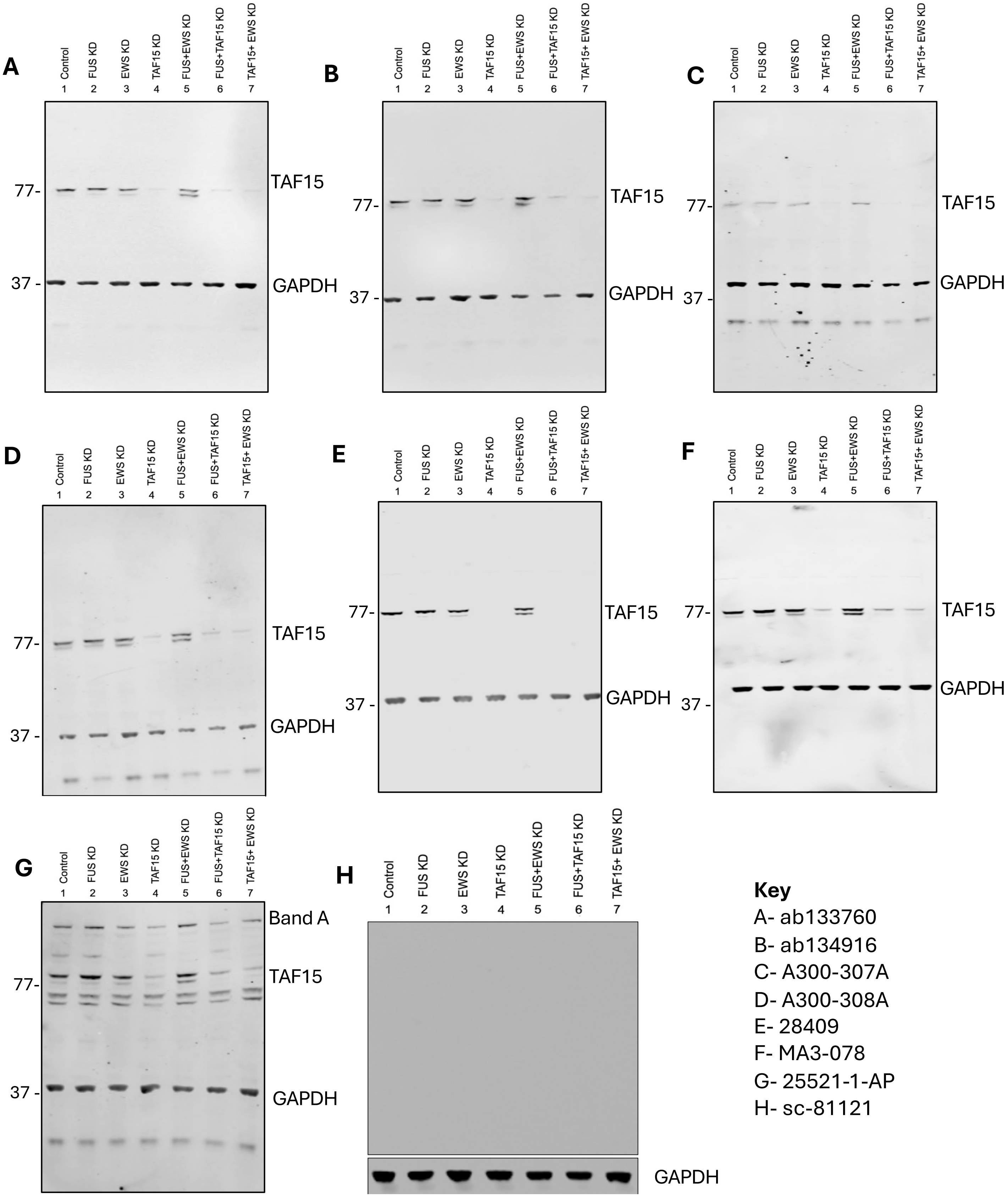


**Figure S9: Anti-TAF15 antibody validation using single/double FET shRNA HeLa knockdowns.** Western blot validation of seven commercially available anti-TAF15 antibodies showed a main band at ~77 kDa corresponding to TAF15. Most antibodies (A, B, D, E, F & G) had an additional lower band determined to be TAF15. All antibodies show a significant decrease of the main TAF15 band in TAF15 only KD (Lane 4), FUS + TAF15 KD (Lane 6) and EWS + TAF15 KD (Lane 7). The lower band increased in the FUS+EWS KD (Lane 5). All bands were normalised to GAPDH loading control (37 kDa) and ratios of FET single/double knockdowns were then compared to control and significance determined by performing One-Way ANOVA with post-hoc Tukey's multiple comparisons test. (see Figure S8 for quantification). (H) sc-81121 showed no TAF15 signal at a range of concentrations.

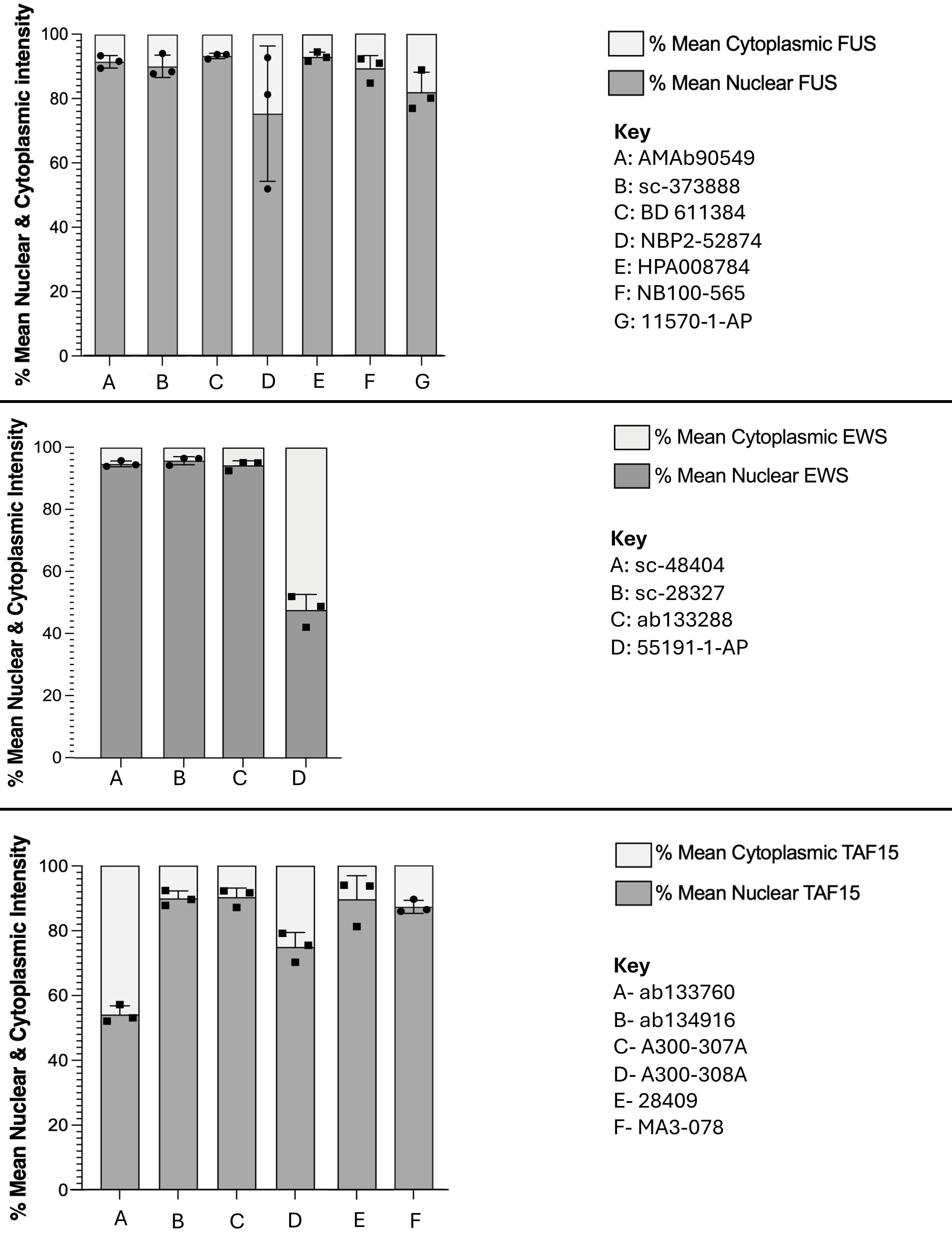


**Figure S10: Quantification of nuclear and cytoplasmic signal for the specific FET antibodies.** Details of quantification methods are in Figure S11. Top panel: graph showing mean percentage of IF signal that is nuclear vs cytoplasmic for the anti-FUS antibodies identified as specific via western blotting. Middle panel: graph showing percentage of IF signal that is nuclear vs cytoplasmic for the anti-EWS antibodies. Bottom panel: graph showing percentage of IF signal that is nuclear vs cytoplasmic for the anti-TAF15 antibodies identified as specific via western blotting.

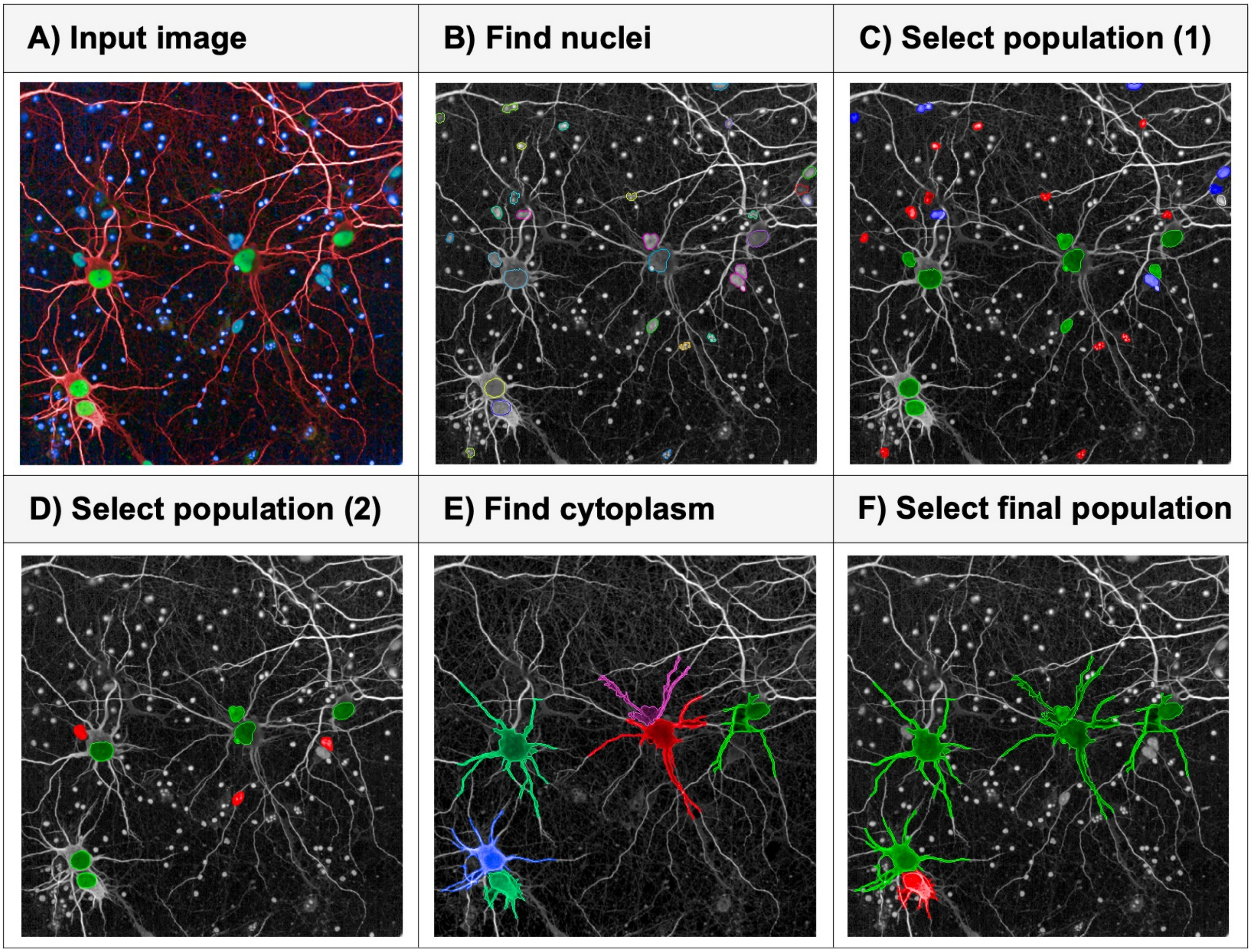


**Figure S11: Overview of 2D analysis pipeline used to quantify FET nuclear and cytoplasmic intensity in rat primary cortical neurons on Harmony.**

A) Input image of selected field of view. B) Nuclei detection using DAPI channel to select Nuclei output population. C) Nuclei filtering step 1, calculated intensity, position, morphology and texture properties from original Nuclei population are used to recognise and stratify new appropriate output populations via linear classification (machine learning) to train software, Neuronal nucleus population (green), Small debris (red), Fuzzy nuclei (blue). D) Nucleus filtering step 2, Neuronal nucleus population is further filtered by using the calculated nucleus area, resulting in a new output population named Main Nuclei. E) The cytoplasm of the Main Nuclei population is detected using the far red (MAP2) channel and intensity, position and morphology properties are calculated from this population. F) Filtering of the final population is carried out by linear classification using the calculated cytoplasmic properties, Final neuronal population (Green) and excluded population (red). The mean FET nuclear and cytoplasmic intensity was then measured from the Final neuronal population. The ratio and percentage mean nuclear and cytoplasmic intensities are then calculated for each FET antibody.
